# MolCodon: A Codon-Based Molecular Language for Interpretable Structural Representation and Similarity Search

**DOI:** 10.64898/2026.05.20.726468

**Authors:** Ehsan Sayyah, Emel Kurul, Huseyin Tunc, Serdar Durdağı

**Author notes:** Corresponding author: (SD).

## Abstract

Molecular representation determines which aspects of chemical structure can be learned, compared, and interpreted in computational drug discovery. Existing encodings typically emphasize either compact string description, as in SMILES and SELFIES, or efficient similarity search, as in circular fingerprints, but they may not simultaneously provide deterministic sequence structure, graph-level interpretability, pharmacophore annotation, and high-fidelity molecular reconstruction. Here, we introduce MolCodon, a codon-based molecular language that represents small molecules as deterministic sequences of fixed-width three-character tokens over a five-symbol alphabet, C, N, O, S, and X. Inspired by the triplet organization of the genetic code, MolCodon assigns chemically defined codon families to atoms, bonds, ring and branch topology, fused-ring references, pharmacophore features, bond mobility, charge, and stereochemistry. A deterministic graph traversal with ring-contiguity preservation produces sequences in which chemically meaningful substructures remain locally organized and traceable to the underlying molecular graph. Across around 2,9 million molecules from six commercial screening libraries, MolCodon achieved 98.93% InChIKey-level round-trip fidelity, supporting its use as a high-fidelity sequence representation for drug-like chemistry. MolCodon-derived sparse sequence and trace features further outperformed SELFIES and Group SELFIES across ten QSAR tasks and exceeded classical fingerprint baselines in six out of ten tasks. As an application of the representation, MolCodon BLAST similarity engine decomposes molecular similarity into ring topology, branch context, attachment architecture, and pharmacophore correspondence, enabling interpretable scaffold-hopping searches. In a PARP1 virtual screening study, MolCodon retrieved scaffold-diverse candidates to a known PARP-1 inhibitor Olaparib. Together, these results establish MolCodon as a new molecular representation paradigm that transforms chemical graphs into high-fidelity, interpretable, and alignment-compatible codon sequences, opening a direct path for bioinformatics-inspired analysis of small-molecule chemical space.

The MolCodon encoder, decoder, and BLAST similarity engine are freely available as open-source software at https://github.com/DurdagiLab/MolCodon

## 1. Introduction

Molecular representation is a foundational problem in computational drug discovery. The form in which a molecule is encoded determines which structural features are available to downstream algorithms, how molecular similarity is defined, whether predictions can be interpreted chemically, and whether encoded structures can be reconstructed as valid molecules. Across virtual screening, quantitative structure–activity relationships (QSAR), scaffold hopping, molecular generation, and library annotation, an ideal representation would combine deterministic encoding, high-fidelity reconstruction, explicit graph-topological information, chemically meaningful local annotations, compatibility with sequence-based computation, and interpretability at the level of atoms, bonds, rings, substituents, and pharmacophore features [1, 2].

Existing molecular representations have made substantial contributions to cheminformatics, although each was developed with particular priorities that leave room for complementary approaches. SMILES remains the most widely used line notation because it is compact, human-readable, and broadly supported by chemical software packages [3]. As a traversal-based string representation of a molecular graph, however, SMILES encodes rings, branches, and long-range graph connections through syntactic markers that may separate directly bonded atoms in the resulting sequence [3, 4]. Consequently, similarity measured at the string level does not always correspond closely to similarity at the molecular-graph level. This can make direct substring matching or sequence alignment challenging to interpret as evidence of shared chemical substructure, especially for ring-containing drug-like molecules in which common cyclic motifs may appear as noncontiguous string patterns. SELFIES addresses a different and highly important objective: Ensuring that arbitrary token sequences decode into chemically valid molecules [5]. This property has made SELFIES particularly useful for generative modeling. At the same time, SELFIES was not primarily designed to represent ring topology, bond mobility, pharmacophore context, or substituent architecture as explicit sequence-level features. DeepSMILES improves some aspects of SMILES syntax and reduces certain string-generation errors [6], although graph-neighbor relationships and chemically meaningful substructures can still be distributed nonlocally across the sequence. InChI provides a robust and widely accepted framework for structure standardization and exact chemical identification [7], but its design is less suited to molecular similarity analysis, machine learning (ML), or interpretable substructure comparison.

Molecular fingerprints represent another important category of molecular descriptors. Extended-connectivity and Morgan fingerprints are widely used in ligand-based virtual screening, largely because they support fast Tanimoto similarity searches across large chemical libraries [8, 9]. Their utility stems from the central role of local atomic environments in many cheminformatics applications. Nonetheless, these representations inevitably lose some structural information and are usually not directly reversible. They omit sequential information, capture global ring systems and substituent organization only incompletely, and often make it difficult to identify the specific structural features responsible for a similarity score. As a result, molecules with high fingerprint similarity can still differ substantially in scaffold topology or pharmacophore positioning, while scaffold-hopping candidates with low fingerprint similarity may retain meaningful biological, topological, or functional correspondence. Learned molecular representations, including graph neural networks (GNNs) and molecular language models, have expanded the predictive power of computational chemistry models [10–13]. These approaches can capture complex structure–property relationships from large datasets, but their learned embeddings are often dense and difficult to decompose into chemically interpretable components.

These limitations point to a broader unresolved challenge. A widely used molecular representation language that is human-readable, expressible over a small symbolic alphabet, and simultaneously supports deterministic sequence encoding, high-fidelity molecular reconstruction, explicit in-sequence chemical annotation, ring-contiguous organization of cyclic systems, and decomposable structural interpretation is currently lacking. Such a representation would not merely be another molecular string. It would define a chemically annotated molecular language in which graph-derived features are organized as interpretable sequence elements, allowing small molecules to be analyzed using sequence-inspired computational strategies while retaining a direct connection to their underlying chemical graphs.

Here, we introduce MolCodon, a codon-based molecular language designed to address this gap. MolCodon represents small molecules as deterministic sequences of fixed-width, three-character codons drawn from a five-symbol alphabet: C, N, O, S, and X. The design is inspired by the triplet organization of the genetic code, but the codon vocabulary is chemically defined. Dedicated codon families encode atom identity, bond order, ring and branch topology, fused-ring references, formal charge, stereochemical state, bond mobility, and pharmacophore annotations. Each emitted codon therefore carries an explicit chemical role rather than functioning as an arbitrary character, traversal artifact, or opaque learned token.

A central design principle of MolCodon is that molecular structure can be expressed through a compact, fixed-width symbolic language rather than through heterogeneous chemical strings or opaque numerical vectors. By restricting the representation to three-character codons drawn from only five symbols, C, N, O, S, and X, MolCodon creates a finite codon space that is small enough to remain computationally tractable, yet sufficiently expressive to encode chemically distinct structural instructions. This design is important for three reasons. First, fixed-width codons remove tokenization ambiguity and make every emitted unit directly comparable across molecules. Second, the small alphabet makes molecular sequences compatible with mature bioinformatics algorithms developed for biological sequence analysis, including local alignment and profile-based search. Third, assigning each codon family to a defined chemical role preserves interpretability: atoms, bonds, ring and branch topology, pharmacophore annotations, bond mobility, charge, and stereochemistry are represented as explicit sequence elements rather than being hidden in hashed bits or latent embeddings. Thus, the five-symbol triplet architecture is not merely a stylistic analogy to the genetic code. It is the mechanism that allows MolCodon to combine compactness, determinism, chemical readability, and alignment-compatible molecular comparison in a single representation.

MolCodon is intended as a general-purpose molecular representation rather than a method restricted to a single downstream task. Its fixed-width codon structure supports sparse sequence features, codon n-gram statistics, trace-derived descriptors, and bioinformatics-inspired comparison strategies. At the same time, its chemically assigned vocabulary preserves information that is often implicit in line notations or lost in hashed fingerprints. Thus, MolCodon provides a framework in which molecular topology, local chemical context, pharmacophore organization, and substituent architecture can be encoded, reconstructed, compared, and interpreted within a unified sequence-based representation.

As one application of this representation, we also developed MolCodon BLAST, an interpretable molecular similarity engine. Rather than relying on direct global alignment of codon strings, MolCodon BLAST decomposes molecular similarity into chemically meaningful components: ring topology, branch context, recursive attachment architecture, and pharmacophore correspondence. These component scores are integrated into an overall similarity value accompanied by a structural explanation. This design allows MolCodon BLAST to identify relationships that may be obscured by conventional fingerprint similarity, including scaffold-hopping candidates that preserve functional or topological correspondence despite low Tanimoto similarity.

We evaluate MolCodon across three complementary levels. First, we assess representational fidelity through a large-scale encode–decode benchmark of 2,898,663 molecules from six commercial screening libraries. This benchmark tests whether MolCodon can reconstruct drug-like molecular structures at scale and identifies the specific structural layers at which failures occur. Second, we evaluate MolCodon-derived sparse sequence and trace features on ten quantitative structure-activity relationships (QSAR) benchmark problems spanning six regression and four classification tasks, using MoleculeNet- and ChEMBL-derived datasets [14, 15]. This analysis tests whether the codon vocabulary captures chemically meaningful information beyond molecular reconstruction. Third, we demonstrate the practical utility of MolCodon-guided similarity analysis in a PARP1 virtual screening and repurposing workflow, followed by physics-based validation [16–23].

Together, these studies establish MolCodon as a new molecular representation paradigm: A deterministic, reconstructable, chemically annotated codon language that converts molecular graphs into interpretable sequence objects. By preserving graph-level chemical meaning while enabling sequence-based analysis, MolCodon creates a direct conceptual interface between cheminformatics and bioinformatics-inspired computation for molecular similarity, scaffold exploration, QSAR modeling, and broader analysis of small-molecule chemical space.

## 2. Materials and Methods

### 2.1 The MolCodon Encoding Scheme

MolCodon represents small molecules using fixed-length, three-symbol codons drawn from a five-symbol alphabet, C, N, O, S, X. This design yields 5^3^ = 125 possible codons. Distinct codon classes are assigned to chemically meaningful features, including atom types, bond orders, ring-topology markers, branching patterns, fused-ring junctions, pharmacophore labels, bond-mobility descriptors, and stereochemical configurations. The resulting molecular strings are conceptually analogous to triplet-based biological sequence encodings, such as the genetic codon system. Because MolCodon encodes molecular structures as ordered symbolic sequences, established sequence-analysis algorithms, such as BLAST-like similarity searches, can be repurposed for chemical informatics applications.

#### 2.1.1 Codon Alphabet and Families

The codon assignments implemented in the current MolCodon encoder are summarized in Table 1. Ten heavy-atom element types are supported: carbon (CCC), nitrogen (CCN), oxygen (CCO), sulfur (CCS), fluorine (CNC), chlorine (CNN), bromine (CNO), iodine (CNS), phosphorus (COC), and boron (CON). Bond order is represented by the NCC codon family, which distinguishes single (NCC), double (NCN), triple (NCO), and aromatic (NCS) bonds. Ring topology is encoded using eight indexed ring-open/ring-close codon pairs, ranging from NNO/NNS to ONN/SCN, allowing up to eight ring connections to be tracked simultaneously. Substituent branching is represented by four branch-open/branch-close codon pairs, from NNC/NNN to NSC/NSN. For fused polycyclic systems, shared atoms are represented without duplication by a fusion-marker construct consisting of OCC, followed by a ring-reference codon and a position codon. Additional atom-level annotation codons encode formal charge states, ranging from +2 (CXS) to -2 (CXX), stereochemical configuration, with SXN denoting R and SXO denoting S, and pharmacophore-related features, including hydrogen-bond acceptors (OXN) and polar hydrogens (OXO). Bond-level annotations capture configurational and conformational properties, including E/Z geometry, with SOX for E and SNX for Z, as well as rotational mobility, with NXC denoting rotatable bonds, NCX non-rotatable bonds, NXS ring-constrained bonds, and NXO aromatic-locked bonds. Sequence boundaries are specified by the start codon SCC and the terminal codon SSS.

**Table 1.** Fusion-reference codons are used to identify completed ring slots during the encoding of fused ring systems and include SOC, SON, SOO, SOS, SSC, SSN, and SSO. Position codons specify intra-ring locations and comprise COO, COS, CSC, CSN, CSO, CSS, OOC, OON, OOO, OOS, SCO, SCS, SNC, SNN, SNO, and SNS. To maintain internal consistency and avoid discrepancies between software components, all downstream analysis programs retrieve codon definitions directly from the MolCodon encoder rather than relying on independently maintained assignment tables.

| Family | Codons | Role | Capacity |
| --- | --- | --- | --- |
| Atom | CCC CCN CCO CCS<br>CNC CNN CNO CNS<br>COC CON | C / N / O / S / F / Cl / Br / I<br>/ P / B | 10 elements |
| Bond type | NCC NCN NCO NCS | Single / double / triple /<br>aromatic | 4 bond types |
| Ring<br>open/close | NNO-ONN / NNS-SCN<br>(8 pairs) | Ring block delimiters with slot<br>index | 8 simultaneous<br>open rings |
| Branch<br>open/close | NNC NOC NOS NSC /<br>NNN NON NOO NSN | Substituent block delimiters | 4 branch depth<br>levels |
| Fusion marker | OCC + RING_REF +<br>POSITION | Shared atom in fused ring (no<br>re-emission) | 7 ref rings × 16<br>positions |
| Charge<br>annotation | CCX CXN CXS CXO<br>CXX | Formal charge 0, +1, +2, -1, -2 | 5 charge states |
| Stereo<br>annotation | SXN SXO | R / S configuration | 2 states |
| Pharmacophore | OXN OXO | H-bond acceptor / polar<br>hydrogen (per H) | Per-atom |
| Bond mobility | NXC NCX NXS NXO | Rotatable / non-rot. / ring /<br>aromatic | 4 states |
| E/Z stereo | SOX SNX | E / Z double-bond geometry | 2 states |
| Control | SCC SSS | Sequence start / end | — |

### 2.2 Deterministic Encoder

Through five consecutive stages, the encoder (*encode*(*smiles*)) transforms a SMILES string into a canonical MolCodon codon list. SMILES-to-graph parsing, Kekulé bond assignment, canonical atom ranking, and stereochemistry perception are the only uses for RDKit, all traversal and emission logic is custom.

#### 2.2.1 Pre-processing

The encoding pipeline begins by converting the input SMILES string into a molecular graph using RDKit. Prior to graph traversal, the structure undergoes two preprocessing steps. The first is Kekulization. In RDKit, aromatic systems are represented using delocalized aromatic bond types; for MolCodon encoding, these bonds must be converted into an explicit Kekulé form with alternating single and double bonds before bond codons can be assigned. Importantly, the aromatic status of each bond is recorded in a lookup set before Kekulization is performed. This preserves information about which bonds were aromatic in the original RDKit representation. The step is required because such bonds must later receive the aromatic-locked mobility annotation, NXO, even though they are represented as single or double bonds after Kekulization. Without this pre-Kekulization record, the encoder would not be able to reliably distinguish originally aromatic ring bonds from ordinary non-aromatic ring bonds in the Kekulized molecular graph.

The second preprocessing step is stereochemical perception. RDKit is used to assign Cahn–Ingold–Prelog (CIP) priority descriptors on the Kekulized molecular graph, identifying tetrahedral stereocenters as R or S and stereochemically defined double bonds as E or Z. These stereochemical labels are stored during preprocessing and subsequently used as atom- or bond-level annotation codons during the MolCodon generation step.

After these preprocessing steps, a canonical atom-rank array is generated for the molecule using RDKit’s canonical ranking algorithm. The resulting array assigns each atom a deterministic integer rank derived from its local and extended chemical environment. The canonical rank does not by itself define the graph traversal order. Instead, it is used only when multiple candidate atoms remain equivalent after applying chemical criteria such as element identity, bond context, and aromatic character. In such cases, the atom with the appropriate canonical rank is selected as a deterministic tiebreaker. Incorporating canonical atom ranks in this manner ensures that chemically equivalent input SMILES strings yield the same MolCodon codon sequence, thereby supporting deterministic and representation-independent encoding.

#### 2.2.2 Start Atom Selection

Atom priority is a fixed chemical ordering (C=0, O=1, N=2, S=3, P=4, halogens 5–8, B=9), aromatic rank is 0 for aromatic atoms and 1 for non-aromatic atoms, and canonical rank is the integer assigned by RDKit canonical ranking. The traversal entry point is the atom with the lowest value of the composite key (atom priority, aromatic rank, canonical rank, and atom index). For every molecular graph, this four-component lexicographic key ensures a distinct, deterministic start atom, generating a single canonical sequence for each molecule.

#### 2.2.3 Graph Traversal and Branch Encoding

Starting from the selected entry atom, the encoder convert the molecular graph using a depth-first strategy. The first thing it checks at each unvisited atom is whether that atom belongs to an unvisited ring. If it does, control is passed immediately to the ring encoder regardless of how the atom was reached, to ensures that ring atoms always form a contiguous block in the output sequence. Non-ring chain atoms are handled straightforwardly to see the bond leading to the atom is emitted first, followed by the atom codon and its annotations.

When an atom has more than one unvisited neighbour, those neighbours are ranked by a composite priority key that extends the atom priority with bond-order rank (aromatic before double, double before triple, triple before single) and uses the bond index as a final tiebreaker. The last neighbour in this ranked list continues the main chain directly. Every other neighbour is enclosed in a branch block. The encoder emits a branch-open codon, recurses into that subtree, and then emits the matching branch-close codon before moving on. Four branch-open/close codon pairs are available, selected by the current branch counter modulo. These four slots refer to available codon labels rather than to any hard limit on nesting depth, so the scheme accommodates molecules with arbitrarily many substituents.

#### 2.2.4 Ring Encoding

When a ring atom is reached during traversal, the encoder allocates a slot from a reusable pool of eight ring-label slots. A ring-open codon for that slot is emitted, and the ring atoms are then visited in a fixed deterministic order starting from the entry atom, with each successive atom chosen by the same neighbour priority key used elsewhere. The bond between consecutive ring atoms is emitted before each atom codon. Once the final atom is processed, the ring-closing bond is emitted and the matching ring-close codon ends the block. The slot is returned to the pool so it can be reused for subsequent rings.

The more important aspect of ring encoding is how substituents and fused-ring connections are handled when they are discovered on ring atoms mid-traversal. Rather than expanding them immediately and breaking the contiguous ring block, the encoder places a label codon at the point of discovery and pushes the actual expansion onto a deferred list. Only after the ring-close codon is emitted are the deferred items processed, in the order they were found. Branch items are expanded as normal branch blocks at that point, and fused-ring connections trigger a recursive ring-encoding call preceded by the appropriate reference and position codons. This approach is what enforces the ring-contiguity property that makes substring matching of ring substructures possible.

#### 2.2.5 Fused Ring Encoding

In fused polycyclic systems, an atom may be shared by two or more rings and may therefore be encountered again during traversal of a subsequent ring. In this case, the encoder does not re-emit the complete atom record, including its annotations, because this information has already been emitted at the atom’s first occurrence. Instead, the shared atom is represented by a compact three-codon fusion signature consisting of the fusion marker OCC, a ring-reference codon identifying the previously completed ring that contains the shared atom, and a position codon specifying the atom’s index within that ring. The corresponding element codon is then emitted immediately after this fusion signature. This representation provides the decoder with sufficient information to retrieve the previously constructed atom and establish the appropriate bond, while avoiding duplication of atom-level structural and annotation data. For deferred fused-ring expansion, a corresponding three-codon header is emitted before the recursive ring-encoding call after closure of the current ring block. This header identifies the parent-ring atom from which the fused connection originates and specifies its intra-ring position, ensuring that the inter-ring bond is reconstructed at the correct location during decoding.

### 2.2.6 Annotation Emission

After each atom codon, the encoder emits zero or more atom-level annotation codons. A formal-charge codon is emitted for every atom: CCX for neutral atoms, CXN for +1, CXS for charges ≥+2, CXO for -1, and CXX for charges ≤-2, respectively. If the atom has an assigned CIP stereochemical descriptor, the corresponding stereochemical codon is appended, with SXN denoting R and SXO denoting S. Pharmacophore-related annotations are also encoded at the atom level. In the current implementation, nitrogen and oxygen atoms receive the hydrogen-bond acceptor codon OXN, and each implicit polar hydrogen attached to the atom is represented by one OXO codon, providing an explicit sequence-level annotation of hydrogen-bond donor capacity. Atoms encountered a second time as part of a fused-ring reference are not re-annotated, because their atom-level information has already been emitted at the first occurrence. Bond codons are similarly followed by bond-level annotation codons. Ring bonds that were aromatic before Kekulization receive the aromatic-locked mobility codon NXO, whereas other ring bonds receive the ring-constrained codon NXS. Acyclic single bonds are annotated as rotatable (NXC) when both endpoints are nonterminal atoms, and as nonrotatable (NCX) when either endpoint is terminal. Acyclic double and triple bonds are always assigned the nonrotatable codon NCX. For stereochemically defined double bonds, the configurational codon is emitted after the mobility annotation, with SOX denoting E geometry and SNX denoting Z geometry.

### 2.3 Stack-Based Decoder

The decoder reconstructs a molecular graph from a whitespace-delimited MolCodon sequence by consuming the tokens from left to right. Its control flow is intentionally parallel to that of the encoder. Whereas the encoder uses recursive graph traversal to emit codons, the decoder uses recursive descent parsing to consume those codons and rebuild the molecular graph.

After the start codon SCC is read, the decoder examines the next token to determine the initial parsing mode. If the token is a ring-open codon, ring parsing is initiated. If it is an atom codon, the corresponding atom is added to the graph and chain parsing proceeds from that atom. During parsing, each bond codon is stored temporarily as a pending bond, together with any associated bond-level annotation codons. The pending bond is applied only after the next atom or ring attachment point has been resolved. Within ring blocks, the decoder follows the same deferred-expansion logic used during encoding. Branch-open and ring-open codons encountered inside a ring are not expanded immediately. Instead, they are placed in a deferred list together with the current atom, which defines the attachment context. Once the corresponding ring-close codon is reached, deferred branches and fused-ring extensions are expanded in the order in which they were recorded.

Fusion signatures are handled by a dedicated parsing routine. When the decoder encounters the fusion marker OCC, it reads the subsequent ring-reference and position codons, identifies the corresponding atom in the completed-ring table, and returns the index of that existing atom rather than creating a new graph node. The following element codon is then checked against the stored atom type as a consistency check.

Stereochemistry is treated differently from most other annotations. In RDKit, the internal chiral tag assigned to an atom depends on the order in which its neighbours are added to the molecular graph. Because the decoder does not necessarily reconstruct atoms in the same neighbour-ordering context used during encoding, directly converting an encoded R or S descriptor into an RDKit chiral tag can introduce systematic errors. For this reason, the CIP stereochemical descriptor is preserved as a custom atom property, MolCodon-Stereo, rather than being converted during graph reconstruction. This preserves the stereochemical annotation in an unambiguous form and allows downstream machine-learning workflows to access it directly.

### 2.4 Trace Annotation

When re-encoding a molecule, the trace module generates an annotation table token by token. Each trace entry contains the codon, its structural role (start, end, atom, bond, ring_open, ring_close, branch_open, branch_close, fusion, fusion_ref, fusion_pos, atom_annotation, bond_annotation), the molecular atom index or bond index it represents, the component type (ring, branch, or backbone), and a component identifier. Component boundaries are preserved with a push/pop stack that accurately mimics the encoder’s delayed paradigm. The created TraceToken list serves as the input for all downstream scoring processes.

### 2.5 MolCodon BLAST Similarity Engine

Because canonical traversal orders are molecule-dependent, identical substructures do not necessarily appear at corresponding sequence positions in different MolCodon strings. Consequently, direct application of conventional sequence-alignment methods, such as Smith–Waterman or BLAST, can give misleading estimates of molecular similarity. MolCodon BLAST therefore does not rely on a single positional alignment score. Instead, molecular similarity is decomposed into five complementary scoring terms, each specifically designed to capture a distinct aspect of structural relatedness. These terms are then integrated into a single overall similarity score that remains interpretable at the level of both the full molecule and its contributing structural features.

#### 2.5.1 Component Extraction

Before any scoring take place, each molecule’s codon sequence is partitioned into structural components using the trace annotation produced during encoding.

The trace records the role and component membership of every token, making it straightforward to identify where each ring, branch, and backbone segment begins and ends.

Ring components are the codon blocks enclosed by matching ring-open and ring-close tokens. Once extracted, each ring block is reduced to a topology-only signature by two normalization steps. First, any Kekulized bond codon that is immediately followed by the aromatic-locked annotation NXO is replaced by the canonical aromatic bond codon NCS, removing differences that arise purely from Kekulization.

Second, all non-topological tokens are stripped, leaving only atom and bond codons. The cleaned sequence is then rotated, all possible shifts of the atom-bond pair sequence are enumerated and the lexicographically smallest rotation is chosen as the canonical ring signature. This makes ring comparison independent of which atom happened to be the traversal entry point.

Branch components are the codon blocks enclosed by matching branch-open and branch-close tokens. Each branch is annotated with its parent component type, either ring or backbone, and with the canonical signature of the parent ring if one exists. This context annotation is what allows the scoring step to distinguish a hydroxyl group on an aromatic ring from a hydroxyl group on an aliphatic chain. Branch records also store any destination ring reached at the branch terminus, as well as any child branches nested within them, building up a recursive fragment-tree structure used in attachment scoring.

Tokens that fall outside all ring and branch blocks form the backbone. Separately, global multiset counters are maintained over all bond-type codons and all pharmacophore annotation codons in the full sequence, used by the bond and pharmacophore scoring steps, respectively.

#### 2.5.2 Similarity Scoring

Following component extraction, four similarity scores are calculated, each describing a distinct aspect of molecular correspondence. For each component, precision and recall are defined from the number of matched features relative to the total number of features in the hit and reference molecules, respectively. The component score is reported as an F1 value scaled to 0–100:

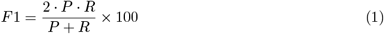

where P is precision, and R is recall. The Ring F1 score is calculated from topology-only ring signatures. Each reference ring is matched to a hit ring with the same canonical ring signature. When several hit rings have the same topology, the match is assigned to the candidate whose attached branch signatures give the highest Jaccard overlap with those of the reference ring. The Branch F1 score is based on context-aware branch keys. Each key combines the parent component type with the branch codon sequence, so that ring-attached branches are compared only with other ring-attached branches, and backbone branches only with other backbone branches. If multiple branches share the same key, ties are resolved using parent-signature identity, recursive fragment-tree similarity, and whether the branch is attached to a ring pair that was already matched during ring scoring. The Attachment F1 score uses graded rather than binary matching. Each attachment is represented by a hierarchical fragment signature that includes its local topology, child fragment trees, and, when applicable, the destination ring signature. Similarity between two attachment trees is computed recursively. If the local signatures differ, the similarity is zero. If they match, a base similarity contribution is assigned, and the remaining score is distributed between child-tree similarity, computed by greedy maximum-weight matching, and agreement of the destination ring. Precision and recall are then accumulated from these continuous similarity values. This allows partial credit for molecules that preserve a common scaffold but differ in substituent composition or extension. The Pharmacophore F1 score is calculated as a multiset F1 over pharmacophore feature–context pairs. The pharmacophore feature specifies the annotation type, and the context is obtained from the trace annotation of the corresponding atom. The four component scores are combined into a single overall MolCodon BLAST similarity score. The attachment score receives double weight to reflect the high informativeness of substituent architecture for structure-activity analysis:

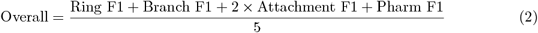

An additional backbone similarity score is computed as a 3-gram Jaccard similarity over backbone tokens and reported alongside the overall score for reference, but it does not contribute to candidate ranking. The Tanimoto coefficient based on Morgan fingerprints (radius 2, 2048 bits) is also computed independently using RDKit and reported for comparative purposes.

### 2.6 QSAR Benchmark

This benchmark was designed to evaluate MolCodon as a supervised QSAR feature source while preserving its primary role as a chemically annotated molecular language for structural similarity analysis. The experiments were organized as task-wise comparisons rather than as a single aggregate leaderboard, because each benchmark task differs in molecular size range, response type, class balance, and chemical diversity. All representation families were therefore processed under the same curation, splitting, model-selection, and evaluation workflow. This design allowed performance differences to be attributed to the molecular representation rather than to differences in preprocessing or model selection.

Let *D*_*t*_ denote benchmark task *t*, where each entry contains a SMILES string *s*_*i*_ and a continuous response or binary label *y*_*i*_.

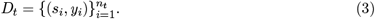

Ten curated QSAR tasks were evaluated, comprising six regression tasks and four binary classification tasks (Table 2). QM7 atomization energy was taken from the MoleculeNet benchmark panel, together with ESOL aqueous solubility, FreeSolv hydration free energy, Lipophilicity, BBBP blood-brain barrier permeability, and the Tox21 NR-AR and SR-p53 toxicity tasks [14]. Tox21 was represented by two tasks rather than the full multitask panel to avoid allowing one benchmark family to dominate the analysis. BCL2 regression was derived from the FDA-approved BCL2 inhibitor repurposing dataset reported by Sayyah et al. [24]. The HIV regression and classification tasks were constructed from the Li et al. [25] HIV-1 protease inhibitor dataset, and were not originally included as official MoleculeNet tasks.

**Table 2:**
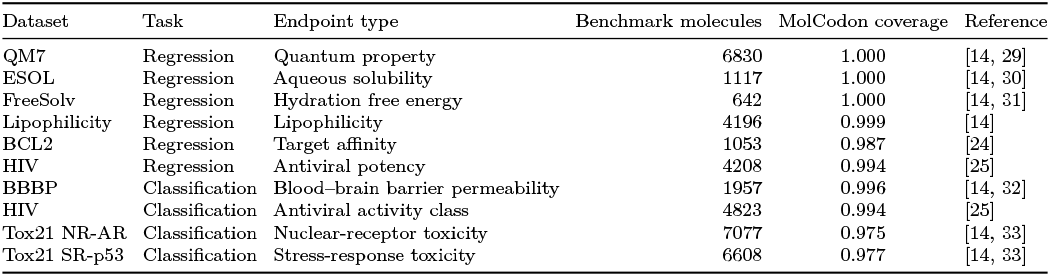
Benchmark datasets and task characteristics.

| Dataset | Task | Endpoint type | Benchmark molecules | MolCodon coverage | Reference |
| --- | --- | --- | --- | --- | --- |
| QM7 | Regression | Quantum property | 6830 | 1.000 | [14, 29] |
| ESOL | Regression | Aqueous solubility | 1117 | 1.000 | [14, 30] |
| FreeSolv | Regression | Hydration free energy | 642 | 1.000 | [14, 31] |
| Lipophilicity | Regression | Lipophilicity | 4196 | 0.999 | [14] |
| BCL2 | Regression | Target affinity | 1053 | 0.987 | [24] |
| HIV | Regression | Antiviral potency | 4208 | 0.994 | [25] |
| BBBP | Classification | Blood–brain barrier permeability | 1957 | 0.996 | [14, 32] |
| HIV | Classification | Antiviral activity class | 4823 | 0.994 | [25] |
| Tox21 NR-AR | Classification | Nuclear-receptor toxicity | 7077 | 0.975 | [14, 33] |
| Tox21 SR-p53 | Classification | Stress-response toxicity | 6608 | 0.977 | [14, 33] |

All molecules were parsed with RDKit and converted to canonical SMILES before feature generation [26]. Invalid structures were removed. Duplicate canonical structures were collapsed before data splitting to reduce train–test leakage, a known concern in QSAR validation when the same molecular structure appears in multiple records. For regression tasks, replicate response values were aggregated using the median,

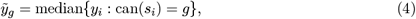

where can(*s*_*i*_) is the canonical SMILES of molecule *i*, and *g* indexes a duplicated canonical structure. Summary statistics for collapsed duplicate groups were retained for auditability. For classification tasks, duplicate molecules were kept only when labels were concordant, and duplicate groups with conflicting labels were removed. All representation families were evaluated on the same final molecule set after RDKit validity, duplicate handling, and MolCodon encodability filtering.

Regression tasks were split using quantile-stratified random 80:20 trainingset–testset partitions. Classification tasks were split using label-stratified random 80:20 partitions. Hyperparameter optimization and model selection were performed by five-fold cross-validation within the training partition only, following the common QSAR practice of separating internal model selection from external held-out evaluation [27]. The independent test partition was not used during hyperparameter tuning, representation selection, or rank aggregation, and was kept for final supporting evaluation.

Table 2 summarizes the final matched benchmark set used for representation comparison. The benchmark molecule count refers to the curated molecules retained after RDKit validity checking, duplicate handling, and MolCodon encodability filtering. MolCodon coverage denotes the fraction of curated molecules that were MolCodon-encodable and retained in the matched benchmark set. Detailed raw-row counts, invalid-structure filtering, duplicate handling, and preprocessing audit columns are reported in Supplementary Table S1.

In addition to the primary random-split benchmark, we conducted a secondary robustness analysis using a Bemis–Murcko scaffold split [14, 28]. Molecules were first grouped by scaffold, and the 80:20 train–test partition was then assigned at the scaffold level, ensuring that scaffolds present in the test set were absent from the training set. The same curated molecular datasets, representation classes, candidate learning algorithms, and cross-validation-based model-selection procedure were used as in the random-split analysis. The scaffold-split performance results, including comparative prediction metrics and model-level summaries, are provided in the Supporting Information (see Supplementary Figures S2 and S3).

#### 2.6.1 MolCodon Encodings

For each molecule, MolCodon features were generated from the deterministic codon sequence and its encoder trace. Six MolCodon-derived QSAR representations were evaluated, namely bag-of-codons (MolCodon-BoC), TF–IDF codon n-grams (MolCodon-TF–IDF), trace features (MolCodon-Tr), hybrid sequence–trace features (MolCodon-Hyb), role-aware features (MolCodon-RA), and super-hybrid features (MolCodon-SH). Let 𝒢_*k*_(*c*_*i*_) denote the set of contiguous *k*-codon n-grams extracted from the MolCodon sequence *c*_*i*_. The bag-of-codons vocabulary was learned from training molecules using unigram and bigram codon features. We use the abbreviation BoC rather than BoW because the retained tokens are MolCodon codons rather than natural-language words, although the construction follows the same sparse bag-of-words/vector-space principle used in text classification and in SMILES LINGO term-frequency models for molecular similarity and drug-target prediction [34, 35].

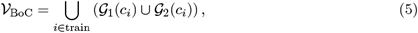

and the corresponding binary feature is

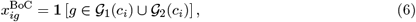

where *g* is a codon n-gram in the learned vocabulary.

The MolCodon-TF–IDF representation expands the vocabulary to one-, two-, and three-codon n-grams, following the same sparse text-vectorization logic commonly used for high-dimensional tokenized inputs and previously adapted to molecular strings through SMILES substring features [34, 35]. With term frequency tf_*ig*_, document frequency df_*g*_, and number of training molecules *N*, the smoothed sublinear TF–IDF weight is

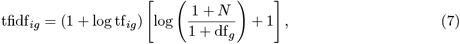

followed by molecule-level *ℓ*_2_ normalization. Vocabularies are learned from the training fold only. Codons or codon n-grams that appear only in validation or test molecules are treated as out-of-vocabulary and contribute zero to the learned sparse vector.

The MolCodon-Tr representation used encoder-trace counters rather than sequence n-grams alone. Let 𝒮_ring_, 𝒮_branch_, 𝒮_attach_, 𝒮_pharm_, 𝒮_bond_, and 𝒮_bb,*k*_ denote vocabularies of normalized ring, branch, attachment, pharmacophore, bond, and backbone *k*-gram signatures. With *n*_*i*_(*s*) as the count of signature *s* for molecule *i*, the trace vector was

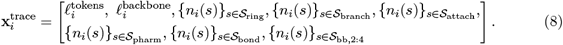

The MolCodon-Hyb representation concatenated sequence and trace channels:

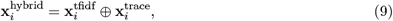

where ⊕ denotes feature concatenation.

The MolCodon-RA representation was designed to prevent chemically distinct codon events from being pooled only because they share the same codon string. Let the trace of molecule *i* be 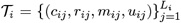, where *c*_*ij*_ is the codon at position *j, r*_*ij*_ is the encoder role, *m*_*ij*_ is the component type (ring, branch, backbone), and *u*_*ij*_ is the component identifier. The role-aware vocabulary was defined by pairing codon n-grams with their trace roles,

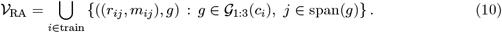

The corresponding role-aware codon feature was the count of each role-conditioned n-gram,

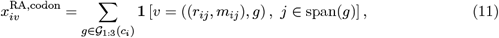

where *v* ∈ 𝒱_RA_. Matched ring and branch blocks were then added as a second role-aware channel. For every component *u*, the opening and closing control codons defined a block *b* = (*m*_*u*_, *h*_*u*_, *a*_*u*_, *d*_*u*_), where *m*_*u*_ is the component type, *h*_*u*_ is the ordered codon signature inside the block, *a*_*u*_ is the codon immediately before entry into the block, and *d*_*u*_ is the codon immediately after exit from the block. The block-context feature vector was

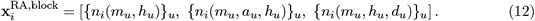

The final MolCodon-RA vector concatenated role-conditioned codon counts and matched block-context counts,

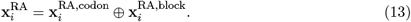

Thus, the same codon can contribute to different coordinates when it appears as an atom, bond, ring delimiter, branch delimiter, pharmacophore annotation, or bond annotation. The MolCodon-SH representation concatenated the TF–IDF sequence channel, MolCodon-RA features, and trace features as

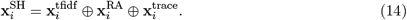

#### 2.6.2 Comptetitor Encodings

SELFIES and Group SELFIES were included as established robust molecular string baselines: SELFIES was designed to guarantee molecular validity under arbitrary strings, whereas Group SELFIES extends this idea with fragment-level group tokens [5, 36]. Both were encoded as token sequences and transformed into sparse bag-of-token and TF–IDF vectors using the same training-fold vocabulary protocol. Fingerprint baselines included ECFP4-2048, FCFP6-2048, RDKit path-2048, and MACCS166. ECFP and FCFP were included as circular fingerprints derived from extended-connectivity fingerprints [8]. MACCS keys were included as a widely used structural-key baseline [37], and RDKit path fingerprints were included as topological-path hashed fingerprints implemented in RDKit [26].

#### 2.6.3 Evaluation Protocol

Candidate estimators included linear, support-vector, tree-based, and gradient-boosting models. Regression candidates were ridge regression, linear support vector regression, random forest [38], LightGBM [39], and XGBoost [40]. Classification candidates were logistic regression, linear support vector machine, log-loss stochastic gradient descent, random forest, LightGBM, and XGBoost. These learners were chosen to span linear sparse models, margin-based models, bagged decision-tree ensembles, and gradient-boosted decision trees, all of which are common baselines for molecular fingerprints and sparse molecular descriptors [41]. Linear models were MaxAbs-scaled, while tree-based and boosting models were fit directly on sparse feature matrices. Hyperparameters were tuned by five-fold cross-validation within the training partition. The complete search grids are provided in Supplementary Table S2.

To avoid selecting a learner from a single endpoint or representation family, global model selection was performed by Borda rank aggregation, a standard voting-based approach for combining rankings across heterogeneous tasks [42–44]. For each task–representation scenario, candidate models were ranked according to the primary cross-validation metric. Regression scenarios were ranked by CV *R*^2^, and classification scenarios were ranked by CV AUROC. The Borda score for model *m* was

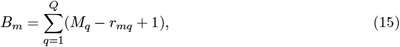

where *Q* is the number of task–representation scenarios, *M*_*q*_ is the number of models available in scenario *q*, and *r*_*mq*_ is the rank of model *m* in that scenario. This analysis was used to evaluate learner robustness across the benchmark, while the main representation-family comparisons report the best cross-validated learner within each representation–task scenario. The full Borda ranking is reported in Supplementary Table S3.

Regression models were evaluated using *R*^2^, RMSE, MAE, Pearson correlation coefficient, and prediction bias. Classification models were evaluated using AUROC, AUPR, accuracy, balanced accuracy, F1 score, and Matthews correlation coefficient. The primary ranking metric was CV *R*^2^ for regression and CV AUROC for classification. Test-set metrics were used as supporting evidence and were not used to choose hyperparameters or representation winners.

All headline comparisons were performed at the task level. For each endpoint, the best MolCodon representation was compared with the best SELFIES-family representation, the best fingerprint representation, and the best non-MolCodon baseline after representation-wise model selection. Differences were reported in the task-primary metric, so regression deltas are differences in CV *R*^2^ and classification deltas are differences in CV AUROC. Because the endpoints differ in size, target type, split protocol, and metric scale, no single pooled performance average was used as the main claim. Complete per-task tables, preprocessing audit columns, scaffold-split diagnostics, and extended flexible-model diagnostics are provided in the Supplementary Information, including the full representation–task grid in Supplementary Table S4.

### 2.7 PARP1 Virtual Screening Protocol

#### 2.7.1 Reference Compound

Olaparib (AZD2281), an FDA-approved PARP1 inhibitor, was used as the primary reference compound [19]. It provides a relevant benchmark for scaffold-hopping analysis because its binding mode within the *NAD*^+^-binding pocket is well characterized and because of its clinical relevance in BRCA-deficient cancers. To broaden the reference set for docking and molecular dynamics (MD) evaluation, two additional high-affinity members of the “-parib” class, talazoparib and saruparib, were included. These compounds were selected on the basis of their low reported IC50 values and distinct binding features, providing complementary reference points for assessing target engagement and the thermodynamic stability of newly identified hits.

#### 2.7.2 Candidate Retrieval

The SPECS 330K diversity library (about 330,000 compounds, [45]) for commercially available novel chemistry and FDA-approved medications (approximately 2,700 compounds, [46]) for repositioning candidates were the two databases that were searched. Following two distinct retrieval operations for each database, the top 50 compounds by MolCodon Overall score were selected from each database. This led to the creation of two candidate sets, SPECS–MolCodon (n=50) and SPECS–FDA (n=50).

#### 2.7.3 Molecular Docking

All candidates from both sets and reference compounds (pH 7.4 ± 0.5, all tautomers) were prepared using LigPrep[20], and they were docked into the PARP1 catalytic site using Glide XP [16, 17] from the Schrodinger Suite (2024-1). The receptor was constructed from PDB entry 7KK4 [19] using the Protein Preparation Wizard [21], with a 20 Å enclosing box centered on the co-crystallized ligand. Glide XP docking scores (kcal/mol) and Glide ligand efficiency (GlideLE = XP score / number of non-hydrogen atoms) were recorded for each molecule. Based on docking score, the top 10 molecules from each dataset were selected for an all-atom molecular dynamics (MD) simulations.

#### 2.7.4 Molecular Dynamics (MD) Simulations

All-atom MD simulations were performed using Desmond [18] with the OPLS4 force field [22] for each of the 10 top-docked compounds per dataset and reference compounds (a total of 23 protein-ligand complexes). Each compound was neutralized with Na^+^/Cl^*−*^ counterions, solvated in an orthorhombic TIP3P water box (10 Å buffer), and equilibrated using the conventional Desmond relaxation procedure. Trajectories were saved every 20 ps throughout production simulations that lasted 20 ns each replica (3 replicates per compound, a total of 69 simulations) at 310 K and 1 atm. Using MM/GBSA [23], the binding free energy was calculated over 100 frames of 20 ns trajectories and averaged across replicas.

## 3. Results

### How MolCodon encodes a molecule

To illustrate the MolCodon encode–decode procedure, we applied the pipeline to two representative drug-like molecules (Figure 1). The encoder traverses the molecular graph in a deterministic depth-first order and emits three-character codons for the structural and annotation features encountered during traversal. Atom codons specify element identity, for example CCC for carbon, CCN for nitrogen, and CCO for oxygen, among the ten supported non-hydrogen (i.e., heavy atom) types. Each atom codon may be followed by atom-level annotations encoding formal charge, CIP stereochemical assignment, hydrogen-bond acceptor status, and the number of polar hydrogens. Bond codons encode bond class, including single (NCC), double (NCN), triple (NCO), and aromatic (NCS) bonds, and may be followed by annotations describing rotatability, E/Z geometry, or ring-constrained character. Branches and rings are represented by paired open and close codons selected from predefined label classes. For fused rings, the encoder additionally writes a fusion marker followed by codons that reference the relevant ring and intra-ring position. Each MolCodon sequence is bounded by the start codon SCC and the stop codon SSS.

**Figure 1:**
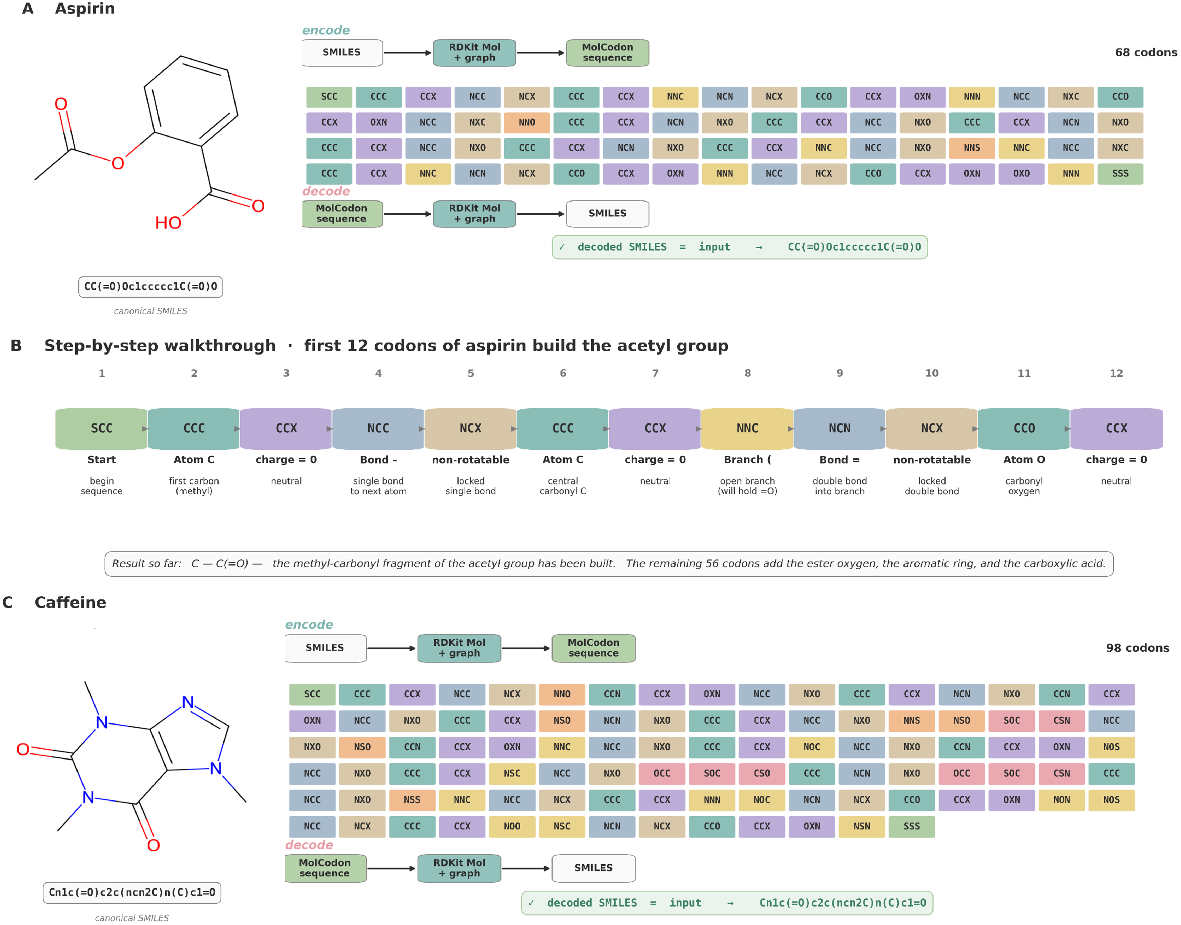
How MolCodon encodes molecules. (A) Aspirin (CC(=O)Oc1ccccc1C(=O)O) yields a 68-codon sequence. Codons are color-coded by structural class. The roundtrip recovers the input canonical SMILES exactly. (B) Step-by-step walkthrough of the first twelve codons of aspirin, which reconstruct the acetyl group. Each codon is annotated with the structural meaning it carries: atom identity, bond type, charge state, rotational mobility, branch delimiter. (C) Caffeine (Cn1c(=O)c2c(ncn2C)n(C)c1=O), a fused bicyclic heteroaromatic, yields a 98-codon sequence using two ring slots and ring-fusion references; the roundtrip again recovers the input exactly.

For aspirin (Figure 1A), the encoder produces a 68-codon sequence. The first 12 codons (Figure 1B) encode the acetyl fragment. The sequence begins with SCC, followed by CCC for the methyl carbon. A single-bond codon (NCC) then connects this atom to the carbonyl carbon, encoded by a second CCC. A branch-open codon (NNC) introduces the carbonyl oxygen branch, which is represented by a double-bond codon (NCN) followed by the oxygen atom codon CCO. The remaining codons encode the ester oxygen, the aromatic benzene ring, and the carboxylic acid substituent. The benzene ring is represented as a contiguous ring block delimited by the slot-0 ring codons NNO and NNS, with aromatic-locked annotations assigned to the corresponding ring bonds.

Caffeine (Figure 1C) provides a more stringent example because it contains a fused bicyclic heteroaromatic scaffold. Its 98-codon MolCodon sequence uses ring slot 0 for the six-membered pyrimidinedione ring and ring slot 1 for the five-membered imidazole ring. Fusion events are encoded using the OCC marker followed by ring-reference and position codons, allowing the decoder to reconnect shared atoms between the two rings without duplicating atom records.

In both examples, decoding the MolCodon sequence reconstructs the original molecular graph, and the resulting canonical SMILES matches the reference canonical SMILES.

### Lossless round-trip benchmark of MolCodon

To evaluate the round-trip fidelity of MolCodon over drug-like chemical space, we performed encode–decode tests on six commercial small-molecule screening libraries: ChemBridge (n = 1,616,862), Enamine (n = 460,160), Specs (n = 328,390), ChemDiv (n = 300,000), OTAVA (n = 160,812), and Life Chemicals (n = 32,439). After de-salting and retention of the largest organic fragment, the combined benchmark comprised 2,898,663 parent structures, providing a large-scale test of molecular representation fidelity across diverse screening collections. Across the full dataset, 2,894,628 molecules encoded successfully, corresponding to 99.86% of the benchmark set. Of the encoded molecules, 2,894,613 decoded to valid RDKit molecular graphs, giving a decoding success rate of 99.999%. Under the strictest graph-identity criterion, 2,870,404 molecules, or 99.03% of the full benchmark, were recovered without detectable structural discrepancy. Per-library and aggregate results are summarized in Figure 2, with the underlying counts reported in Table 3.

**Table 3:** MolCodon roundtrip fidelity across six commercial drug-like screening libraries. Libraries are listed in decreasing size order. Encode and Decode columns report the fraction of input molecules that successfully passed each stage; decode is conditional on encode succeeding. InChIKey is the chemoinformatics-standard lossless roundtrip criterion. Conn. (connectivity) denotes graph identity ignoring stereochemistry. Strict denotes the full graph fingerprint including chirality, double-bond E/Z, ring/aromatic flags, and MolCodon pharmacophore annotations. The Combined row weights each library by its size.

| Library | n | Encode | Decode | InChIKey | Conn. | Strict | Total failures |
| --- | --- | --- | --- | --- | --- | --- | --- |
| ChemBridge | 1,616,862 | 99.76% | 100.00% | 98.43% | 99.33% | 98.70% | 21,056 (1.30%) |
| Enamine | 460,160 | 100.00% | 100.00% | 99.55% | 99.92% | 99.77% | 1,045 (0.23%) |
| Specs | 328,390 | 99.96% | 99.997% | 99.25% | 99.31% | 98.47% | 5,025 (1.53%) |
| ChemDiv | 300,000 | 100.00% | 100.00% | 99.89% | 99.94% | 99.86% | 422 (0.14%) |
| OTAVA | 160,812 | 100.00% | 100.00% | 99.53% | 99.77% | 99.57% | 693 (0.43%) |
| Life Chemicals | 32,439 | 100.00% | 99.99% | 99.99% | 99.99% | 99.94% | 18 (0.06%) |
| Combined | 2,898,663 | 99.86% | 99.999% | 98.93% | 99.51% | 99.03% | 28,259 (0.97%) |

**Figure 2:**
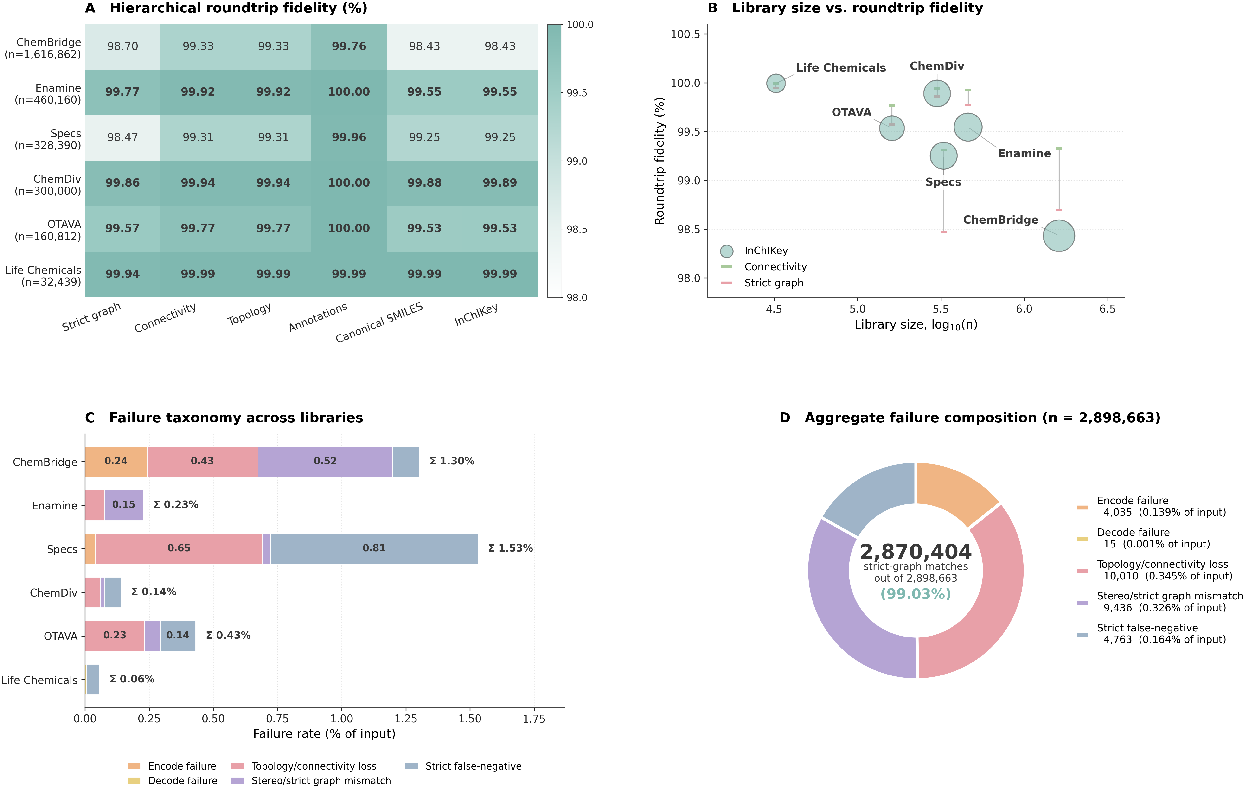
MolCodon roundtrip benchmark across six commercial drug-like screening libraries (combined n = 2,898,663). (A) Heatmap of hierarchical fidelity metrics for each library; cell values are percentages and deeper teal corresponds to higher fidelity within the 98–100% display range. (B) Library size on a logarithmic axis versus fidelity. The bubble marks InChIKey fidelity for each library; the small horizontal dashes mark connectivity (sage) and strict-graph (rose) fidelities for the same library. No systematic dependence of fidelity on library size is observed. (C) Stacked horizontal bars decompose the total failure rate of each library into the five failure classes defined in the text; the cumulative failure rate is annotated at the right of each bar. (D) Aggregate composition of all failures across the combined corpus, expressed as a donut. The center of the donut reports the 2,870,404 / 2,898,663 = 99.03% strict graph match rate.

The libraries were selected to cover a broad range of molecular sizes and structural diversity. ChemBridge and Enamine together contribute more than two million synthetic, medicinal-chemistry-oriented structures and therefore provide most of the statistical weight of the benchmark. Specs adds a higher proportion of natural-product-like scaffolds and less common heteroatom patterns. ChemDiv represents a curated synthetic diversity collection, OTAVA contributes screening-fragment chemistry, and Life Chemicals provides a smaller curated set of drug-like molecules. Together, these sources approximate the range of structures that MolCodon would encounter in practical ligand-discovery workflows rather than reflecting a single vendor-specific chemical space.

In order of increasing permissiveness, we present six complimentary fidelity metrics: strict graph, connectivity, topology, canonical SMILES, MolCodon pharmacophore annotations, and InChIKey. Because the decoded molecule must match the input on atom identity, formal charge, isotope state, bond order, ring and aromatic flags, atom-level CIP stereochemistry, double-bond E/Z stereochemistry, and pharmacophore annotations specific to MolCodon, the strict graph criterion is the most stringent. The chemoinformatics community standard for identifying two structures as identical is called InChIKey identity. The space between these two endpoints is divided by the five intermediate metrics, which allow us to pinpoint any failure to a particular structural layer. In almost all circumstances, a canonical SMILES match implies an InChIKey match, and the hierarchical inclusion is stringent, connectivity, and topology.

We classified failures into five types that have mechanistic significance. When a molecule cannot be transformed into a MolCodon sequence, an encode failure occurs. When a sequence is generated but cannot be rebuilt into an accurate RDKit molecule, this is known as a decode failure. Genuine encoder-decoder corruption of the molecular graph is covered by topology/connectivity loss. Stereo/strict graph mismatch refers to situations in which stereochemistry was lost while the graph topology remained intact. Roundtrips that were chemically successful according to canonical SMILES and InChIKey but that the strict graph fingerprint identified as different are known as strict false-negatives; these are signature artifacts rather than information loss.

Across 2,898,663 molecules, MolCodon achieved 99.86% encode success, 99.999% decode success conditional on successful encoding, 98.93% InChIKey-level lossless roundtrip identity, 99.51% connectivity-level graph identity, and 99.03% strict graph identity. Annotation fidelity, which checks whether MolCodon-encoded pharmacophore features (hydrogen-bond acceptors and polar-hydrogen counts) survived the roundtrip, reached 99.86% on the full corpus and 100.00% on five of the six libraries.

The smallest and largest libraries define the range of observed performance. Life Chemicals (n = 32,439) achieved 99.99% fidelity across all evaluated criteria, whereas ChemBridge (n = 1.6 million) achieved 98.43% InChIKey identity and 98.70% strict graph identity. The four intermediate-sized libraries showed similarly high recovery rates: ChemDiv reached 99.89% InChIKey identity and 99.86% strict graph identity, Enamine 99.55% and 99.77%, OTAVA 99.53% and 99.57%, and Specs 99.25% and 98.47%, respectively. Figure 2A illustrates the expected hierarchy among the fidelity metrics. Connectivity and topology identities were nearly indistinguishable, typically differing by less than 0.01%, and both were consistently higher than strict graph identity by approximately 0.2–1.0%. This difference is mainly attributable to stereochemical perception and annotation sensitivity rather than loss of molecular connectivity. Figure 2B shows fidelity as a function of library size on a logarithmic scale. No systematic relationship was observed between library size and reconstruction fidelity, indicating that the modest variation among libraries is better explained by differences in scaffold composition and chemical complexity than by dataset scale.

The two largest libraries (ChemBridge and Specs) account for the bulk of the absolute failure count and provide the clearest view of the failure mechanisms. ChemBridge contributed 21,056 total failures (1.30% of input), of which 3,904 were encode failures, 6,985 were topology/connectivity losses, and 8,485 were stereo/strict graph mismatches. Specs contributed 5,025 failures (1.53% of input) with a profile dominated by strict false-negatives (2,651 cases) and topology/connectivity losses (2,126 cases). The other four libraries each had less than 0.5% total failure rate.

### Failure mode analysis

Of the 28,259 molecules (0.97% of input) that failed at least one fidelity check, the failures decomposed cleanly into mechanistically distinct populations. These are molecules that could not be encoded at all. They concentrate in libraries that include scaffold-rare chemistry: ChemBridge (3,904 encode failures, 0.24%) and Specs (131 encode failures, 0.04%). The remaining four libraries contained zero encode failures. The dominant root causes are main-group atoms outside MolCodon’s designed alphabet (silicon, selenium, deuterium labels, and trace transition metals such as Ni, Ti, Ge and Fe), and codon alphabet capacity limits reached by molecules with eight or more fused rings or with deeply nested branch labels. Both are designed scope limitations rather than algorithmic defects: the codon alphabet was sized for drug-like organic chemistry (C, N, O, S, halogens, P, B), and the eight-slot ring label LIFO was chosen to keep the codon length tractable.

Only 15 molecules out of 2.9 million produced a sequence that could not be reconstructed by the decoder. Twelve came from ChemBridge’s caged polycyclic fragment, two from Life Chemicals, and one from Specs. These represent the algorithmic boundary at which the deferred-branch label system collides with bridged bicyclic structures — a known scoped-out limitation.

This class contains molecules where the encoded sequence decoded into a chemically different graph: bonds rearranged, ring fusion altered, or atoms relocated. These are the failures that truly represent information loss. ChemBridge (6,985) and Specs (2,126) dominate this class; both libraries contain unusually dense polycyclic scaffolds. ChemDiv (178), OTAVA (373), Enamine (348) and Life Chemicals (0) round out the picture. The structural signature of this class is a high density of fused rings and bridged systems near the edge of MolCodon’s representational capacity.

The connectivity graph survived the roundtrip but at least one stereo feature (a CIP-assigned tetrahedral center or a double-bond E/Z label) was lost, causing InChIKey to disagree. These cases are chemically more subtle: the scaffold and bonding are correct, but the stereochemistry annotation did not round-trip. The class is largest in ChemBridge (8,485) and concentrates in molecules with multiple stereocenters embedded in symmetric ring systems where RDKit’s stereo perception is borderline.

For these molecules, the decoded canonical SMILES and InChIKey were identical to those of the input, whereas the strict graph fingerprint indicated a mismatch. Inspection showed that these cases arose from RDKit’s assignment of STEREOE/STEREOZ labels, which depends on the pair of neighboring atoms selected as stereochemical references. In molecules containing multiple stereocenters or symmetric substituents, canonical-rank tie-breaking can select different reference pairs in the input and decoded graphs. This can invert the reported E/Z descriptor without changing the underlying chemical structure. To verify this interpretation, we applied a canonical-rank-normalized stereo signature, which removed the apparent discrepancies. These cases therefore represent false negatives of the strict graph metric rather than failures of the MolCodon encoding or decoding procedure. The corresponding molecular structures were recovered correctly.

### Interpretation

Combining the genuine failure categories, including encoding failures, decoding failures, topology or connectivity loss, and stereochemical loss, identified 23,496 incorrectly recovered molecules among 2,898,663 input structures. This corresponds to a genuine loss rate of 0.811%. After excluding strict-metric false negatives, which reflect chemically successful round trips affected by metric-specific artifacts, the effective lossless round-trip rate of MolCodon was 99.19% under the strictest chemically interpretable criterion and 98.93% under the more conservative full-InChIKey criterion, including stereochemical information. For practical drug-discovery workflows, including similarity search, scaffold clustering, library annotation, and sequence-based deep learning, operational fidelity is best represented by chemically meaningful identity measures such as full InChIKey agreement. On this basis, MolCodon achieved 98.93% round-trip fidelity across nearly three million curated screening-library structures.

ChemBridge showed a modestly higher failure rate (1.30%), consistent with the breadth of this collection. With 1.6 million structures, ChemBridge samples further into the long tail of medicinal chemistry, including bridged and spirocyclic systems as well as less common heteroatom patterns. Specs also showed an elevated failure rate (1.53%), which is consistent with its enrichment in natural-product-like chemistry; silicon-containing organosilicon fragments alone accounted for 69 encoding failures in this library. By contrast, the four libraries dominated by more conventional drug-like chemistry, ChemDiv, Enamine, OTAVA, and Life Chemicals, each showed total failure rates below 0.5% and InChIKey-level fidelity above 99.5%. These results provide a practical estimate of MolCodon performance within the chemical space for which the representation was primarily designed.

Existing sequence-based molecular representations address different parts of this problem. SMILES remains compact and partly human-readable, but its variable tokenization, traversal-order sensitivity, and chemical syntax differ substantially from biological sequence formats, making direct use with bioinformatics alignment tools nontrivial and often requiring preprocessing or adaptation. SELFIES provides robust syntactic validity, which is valuable for generative modeling, although its tokens are not always aligned with chemically interpretable structural units. MolCodon was designed to provide a complementary representation: A fixed-width, three-character codon system based on a small alphabet, conceptually analogous to biological triplet codes, in which atoms, bonds, rings, branches, stereochemical descriptors, and pharmacophore annotations are encoded explicitly in the sequence. This design makes MolCodon compatible with sequence-analysis frameworks such as Smith–Waterman alignment, BLAST-like search, and profile hidden Markov models, provided that chemically informed scoring schemes are used. At the same time, the encode–decode benchmark shows high round-trip fidelity across large drug-like screening libraries, with identity rates in the 98.93–99.86% range depending on the fidelity criterion applied. To our knowledge, MolCodon is the first fixed-width molecular representation based on a small biological-sequence-like alphabet to be evaluated at this scale while retaining this level of chemically resolved reconstruction fidelity.

### Glossary of metrics and failure classes

The benchmark was evaluated using six fidelity metrics and five failure classes, defined to separate whole-molecule identity from more localized reconstruction errors. The primary lossless round-trip metric is strict graph fidelity. Under this criterion, the decoded molecule must preserve atom identity, formal charge, isotope and radical state, molecular connectivity, bond order, aromatic and ring-membership flags, assigned atom-level CIP stereochemistry, assigned double-bond E/Z geometry, and MolCodon-specific pharmacophore annotations. Because atoms are matched by canonical ranking before comparison, this metric can localize discrepancies to individual atoms, bonds, stereochemical labels, or annotations rather than relying only on whole-molecule string identity.

Connectivity fidelity evaluates whether the decoded molecule preserves the same atom-bond connectivity as the input. It does not require complete agreement in stereochemical or annotation layers. A molecule that passes connectivity fidelity but fails strict graph fidelity therefore retains the molecular scaffold but differs in at least one higher-order feature, such as stereochemical assignment, bond annotation, aromatic perception, or MolCodon-specific atom annotation. This metric is most useful for asking whether the core molecular graph survived the round trip.

Topology fidelity provides a still more permissive graph-level comparison. It tests whether the same atoms remain connected in the same overall graph topology while ignoring bond order and stereochemistry. Failures at this level correspond to chemically important reconstruction errors, such as incorrect branch placement, misplaced fusion bonds, or incomplete ring reassembly. In practice, topology fidelity closely tracks connectivity fidelity because structures that lose graph connectivity are rarely rescued by ignoring bond order.

Annotation fidelity measures preservation of MolCodon-specific atom annotations independently of graph reconstruction. These annotations include features such as hydrogen-bond acceptor status and polar-hydrogen counts. High annotation fidelity indicates that the pharmacophore information encoded directly in the MolCodon sequence remains available for downstream similarity search, library annotation, and machine-learning applications.

Canonical SMILES fidelity compares the RDKit canonical isomeric SMILES generated from the input and decoded molecules. This metric provides a useful string-level consistency check, but it is sensitive to toolkit-dependent canonicalization, aromaticity perception, and stereochemical labeling. Thus, two chemically equivalent structures may occasionally differ at the canonical SMILES level if their perceived aromatic or stereochemical representation changes during reconstruction.

The chemoinformatics community’s standard for identifying two structures as identical is called InChIKey fidelity. InChI is the most chemically meaningful identity check accessible since it uses a tool-independent canonicalization to standardize tautomers, protonation states, and stereochemistry. However, because it does not localize which feature changed when it disagrees, it is less diagnostic than graph metrics.

An encode failure occurs when the input molecule cannot be converted into a MolCodon sequence. The main causes are outside the intended alphabet or capacity of the present implementation, including unsupported elements such as silicon, selenium, transition metals, or deuterium labels, and highly packed polycyclic systems that exceed the available ring-encoding scheme. These cases reflect the defined scope of the current MolCodon alphabet rather than a failure of the traversal algorithm.

A decode failure occurs when a MolCodon sequence is produced but cannot be assembled into a valid RDKit molecular graph. These cases were rare and were concentrated in bridged bicyclic systems near the edge of the ring-labeling logic. They represent cases in which the encoded sequence could not be converted back into a chemically valid graph by the current decoder.

A topology/connectivity loss is recorded when decoding yields a valid molecule, but its connectivity or graph topology differs from that of the input. This class represents genuine information loss because the molecular scaffold was not reconstructed correctly. Examples include incorrect branch attachment, misplaced ring-fusion connections, or failure to recover the original ring system.

A stereo or strict-graph mismatch occurs when the molecular graph is recovered but at least one higher-order feature differs. These cases include changes in double-bond E/Z assignment, atom-level CIP stereochemistry, or other strict-graph features despite preservation of the underlying connectivity. In this class, the graph is correct, but the stereochemical or annotation state is not fully conserved. A strict false negative denotes a metric-level discrepancy rather than a chemically meaningful reconstruction failure. In these cases, the decoded canonical SMILES and InChIKey match the input, but the strict graph fingerprint reports a mismatch. Inspection traced these cases mainly to RDKit STEREOE/STEREOZ labeling, which depends on the neighboring atoms selected as stereochemical references. In molecules with multiple stereocenters or symmetric substituents, canonical-rank tie-breaking can select different reference pairs in the input and decoded graphs, changing the reported E/Z label without changing the chemical structure. These cases were therefore treated as artifacts of the strict comparison metric and were excluded from estimates of genuine MolCodon information loss.

### 3.3 QSAR Benchmarking

#### ML Model Selection

Before comparing representation families, we first examined whether one learner was consistently strong across the full benchmark. The model-selection surface contained 140 task–representation scenarios, defined by crossing the ten endpoints with all evaluated representation views. Within each scenario, models were ranked by the task-primary cross-validation metric *R*^2^ for regression and AUROC for classification. These ranks were then aggregated by Borda scoring [42–44], which avoids directly averaging metrics with different scales.

XGBoost achieved the highest aggregate Borda score and the best mean rank across the full set of benchmark scenarios (Table 4). LightGBM showed comparable performance, with a marginally lower aggregate Borda score but the largest number of first-place rankings. Random forest was the next most consistent learner across representations and tasks. Linear models were generally less competitive, although ridge regression remained useful for selected regression settings. Overall, these results indicate that gradient-boosted tree models and ensemble methods provide the most robust performance for the sparse MolCodon, SELFIES, Group SELFIES, and fingerprint-derived feature spaces evaluated here. Additional model-selection diagnostics, including the Borda-score summary and the mean-rank heatmap stratified by representation family, are reported in Supplementary Figure S1.

**Table 4:** Model-selection ranking. Models were ranked within each task–representation scenario by 5-fold CV *R*^2^ for regression or 5-fold CV AUROC for classification, then aggregated by Borda scoring. Higher Borda scores and lower mean ranks indicate more consistent performance across the benchmark.

| Model | Borda score | Mean rank | Median rank | First-place scenarios | Top-2 scenarios | Scenarios |
| --- | --- | --- | --- | --- | --- | --- |
| XGBoost | 680 | 2.14 | 2.0 | 46 | 94 | 140 |
| LightGBM | 676 | 2.17 | 2.0 | 48 | 91 | 140 |
| Random forest | 630 | 2.50 | 2.0 | 40 | 72 | 140 |
| RBF-SVM | 388 | 4.23 | 4.0 | 3 | 14 | 140 |
| Linear SVM | 297 | 4.88 | 5.0 | 2 | 7 | 140 |
| SGD-SVM | 269 | 5.08 | 5.0 | 1 | 2 | 140 |

**Table 4:** Top 10 compounds by Glide XP docking score per dataset, selected for MD simulation. MM/GBSA values are 3-replica averages from 20 ns simulations.

Reference Compounds
| Compound | MM-GBSA<br>(kcal/mol) | Glide XP<br>(kcal/mol) |
| --- | --- | --- |
| **Olaparib** | **$-100.67 \pm 3.3$** | -15.59 |
| Talazoparib | $-59.13 \pm 3.85$ | -5.3 |
| Saruparib | $-39.35 \pm 8.03$ | -2.88 |

FDA – MolCodon hits
| Compound | MM-GBSA<br>(kcal/mol) | Glide XP<br>(kcal/mol) |
| --- | --- | --- |
| **Oxatomide** | **$-78.5 \pm 4.2$** | -4.8 |
| Fezolinetant | $-60.2 \pm 4.7$ | -5.1 |
| Droperidol | $-60.2 \pm 2.9$ | -4.7 |
| Moxifloxacin | $-57.0 \pm 3.7$ | -4.5 |
| Ataluren | $-48.8 \pm 2.5$ | -4.3 |
| Ciprofloxacin | $-38.8 \pm 4.2$ | -3.9 |
| Rufinamide | $-38.7 \pm 3.4$ | -3.8 |
| Dexketoprofen | $-37.3 \pm 3.2$ | -3.6 |
| Ketoprofen | $-35.4 \pm 3.6$ | -3.5 |
| Cinoxacin | $-33.7 \pm 4.3$ | -3.4 |

SPECS – MolCodon hits
| Compound ID | MM-GBSA<br>(kcal/mol) | Glide XP<br>(kcal/mol) |
| --- | --- | --- |
| **AK-968/40726826** | **$-74.3 \pm 5.8$** | -7.2 |
| AG-690/12949060 | $-56.0 \pm 4.1$ | -6.9 |
| AG-690/12950212 | $-53.6 \pm 3.3$ | -6.7 |
| AT-057/43316532 | $-52.9 \pm 4.8$ | -6.8 |
| AT-057/43314007 | $-51.9 \pm 4.2$ | -6.5 |
| AG-690/12949061 | $-50.8 \pm 5.6$ | -6.6 |
| AG-690/12949049 | $-50.6 \pm 4.5$ | -6.4 |
| AG-690/12949058 | $-47.5 \pm 4.7$ | -6.3 |
| AG-690/12949053 | $-42.5 \pm 2.2$ | -6.1 |
| AT-057/40126606 | $-42.2 \pm 4.2$ | -6.0 |

XGBoost, LightGBM, and random forest formed the top-performing group of learners, whereas linear models showed less consistent performance across representation families. XGBoost was selected for downstream analyses because it achieved the highest aggregate Borda score. However, its margin over LightGBM was small, indicating that the subsequent representation comparisons should be viewed as evaluations under a strong gradient-boosted tree model rather than as evidence that XGBoost is uniquely optimal for these datasets.

XGBoost, LightGBM, and random forest formed the dominant learner tier, while linear models were less stable across representation families. The subsequent representation comparisons therefore report the best cross-validated representation–model pair within each family, with the Borda table retained as a model-selection robustness diagnostic.

#### Regression Tasks

Figure 3 summarizes the six regression tasks. MolCodon-SH was the best MolCodon representation on QM7, ESOL, FreeSolv, Lipophilicity, and HIV regression, while MolCodon-Hyb was best on BCL2 regression. These results indicate that the combined sequence–trace representation was usually more effective than sequence-only MolCodon features.

**Figure 3:**
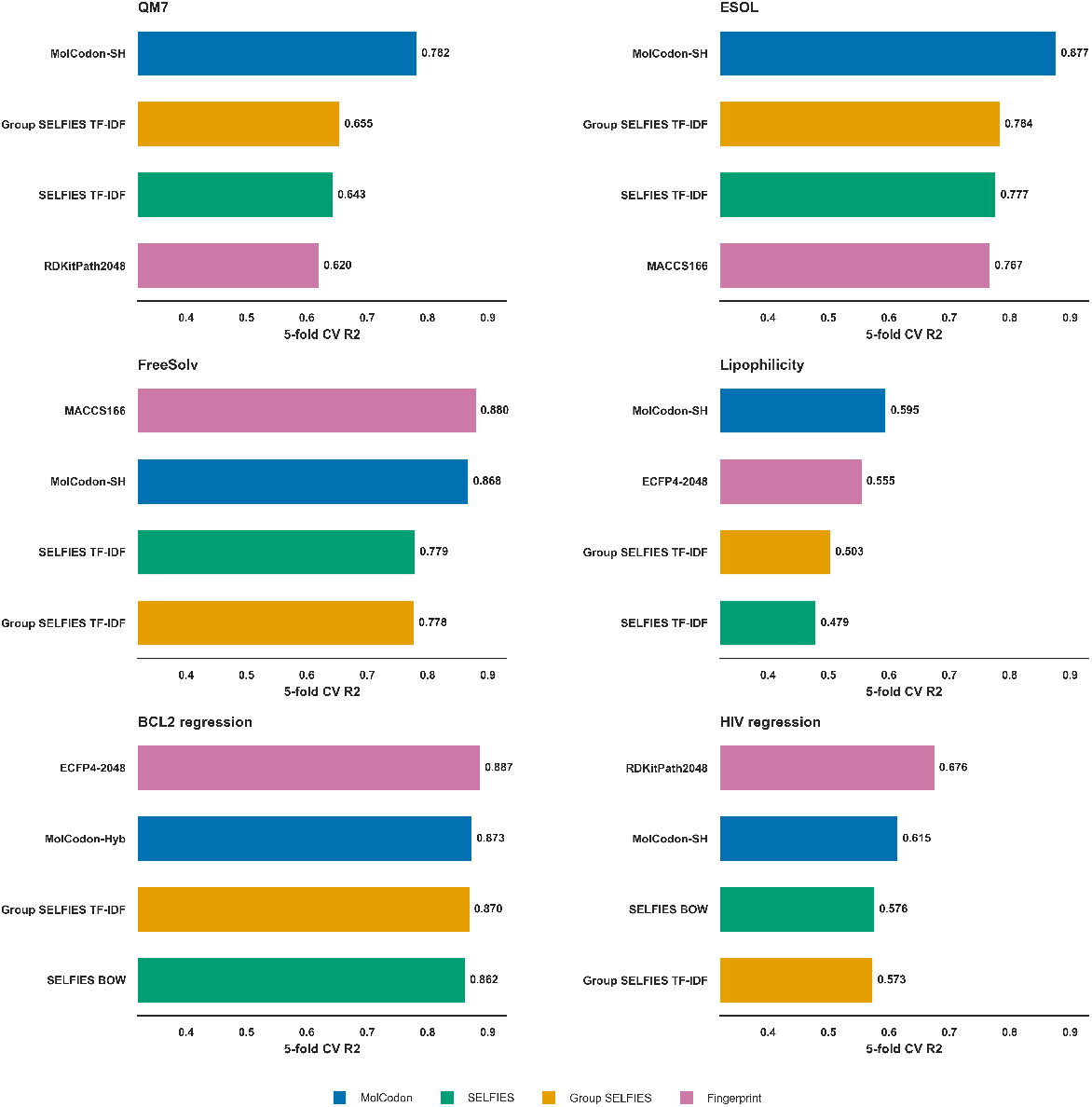
Regression tasks. Best representation within each family on the six regression tasks using the best cross-validated learner for each representation. The primary metric is 5-fold CV *R*^2^.

MolCodon clearly outperformed the other string representations across the regression panel. On QM7, MolCodon-SH reached a CV *R*^2^ of 0.782, compared with 0.655 for Group SELFIES TF–IDF and 0.620 for the best fingerprint baseline. On ESOL, MolCodon-SH reached 0.877 CV *R*^2^, exceeding Group SELFIES TF–IDF (0.784) and MACCS166 (0.767). On Lipophilicity, MolCodon-SH achieved 0.595 CV *R*^2^, above ECFP4-2048 (0.555) and Group SELFIES TF–IDF (0.503).

The regression endpoints also highlight cases in which established fingerprint representations remain highly competitive. For FreeSolv, MACCS166 gave the best performance, with a crossvalidated *R*^2^ of 0.880, slightly above MolCodon-SH at 0.868. For BCL2 regression, ECFP4-2048 was the top representation (*R*^2^=0.887), with MolCodon-Hyb close behind (*R*^2^=0.873). The largest fingerprint advantage was observed for HIV regression, where RDKit path-2048 achieved a cross-validated *R*^2^ of 0.676 compared with 0.615 for MolCodon-SH.

This weaker HIV regression result should be interpreted with the composition of the activity data in mind, because the curated regression table combined IC_50_ and K_*i*_ activity records within a single activity-prediction task.

Taken together, these results show that MolCodon provides a strong sequence-based regression representation, consistently improving over string baselines and approaching the performance of optimized fingerprints, which remained advantageous for a subset of bioactivity endpoints.

The held-out parity plots provide an additional view of regression behavior across the full regression panel (Figure 4). QM7, ESOL, FreeSolv, and BCL2 regression showed close agreement between observed and predicted values, with predictions clustering around the identity line, consistent with the strong performance of the corresponding MolCodon models. Lipophilicity showed a moderate increase in dispersion but retained the overall trend across the test set. HIV regression exhibited the widest spread among the regression endpoints, indicating that this endpoint presents a more challenging activity landscape for sequence-based modeling, although MolCodon still captured the principal structure–response trend.

**Figure 4:**
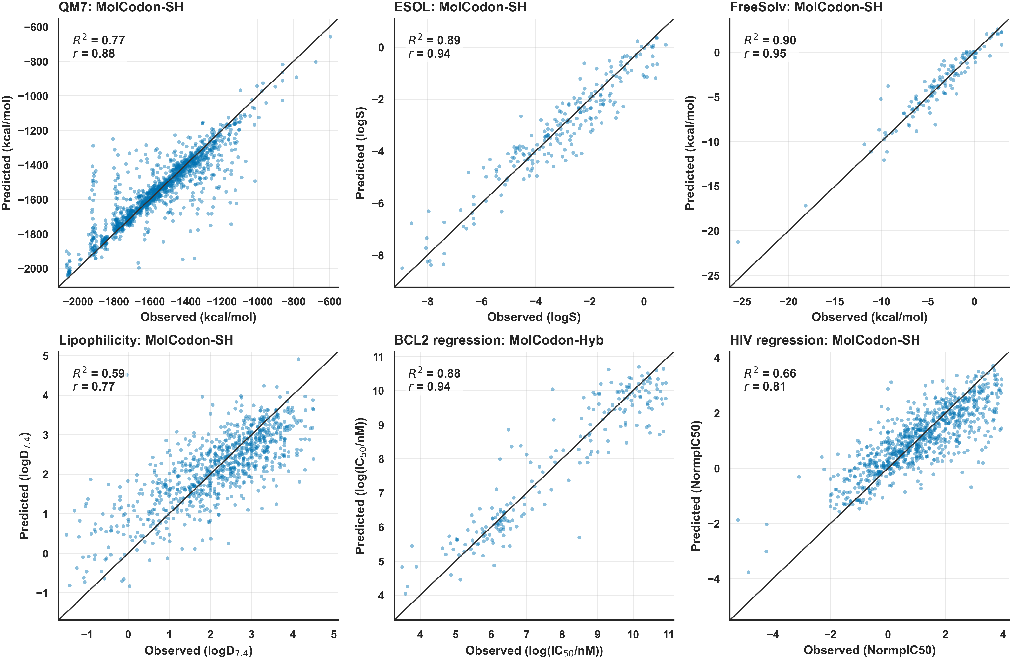
Regression parity diagnostics. Points show held-out test predictions from the MolCodon family-best models for QM7, ESOL, FreeSolv, Lipophilicity, BCL2 regression, and HIV regression. The dashed line indicates perfect agreement between observed and predicted responses.

#### Classification Tasks

Figure 5 summarizes the four classification tasks. MolCodon was competitive across all classification tasks and achieved the best family-level score on two of four tasks. MolCodon-Hyb was best on BBBP, and MolCodon-SH was best on Tox21 SR-p53. Fingerprints achieved the highest AUROC on HIV classification and Tox21 NR-AR.

**Figure 5:**
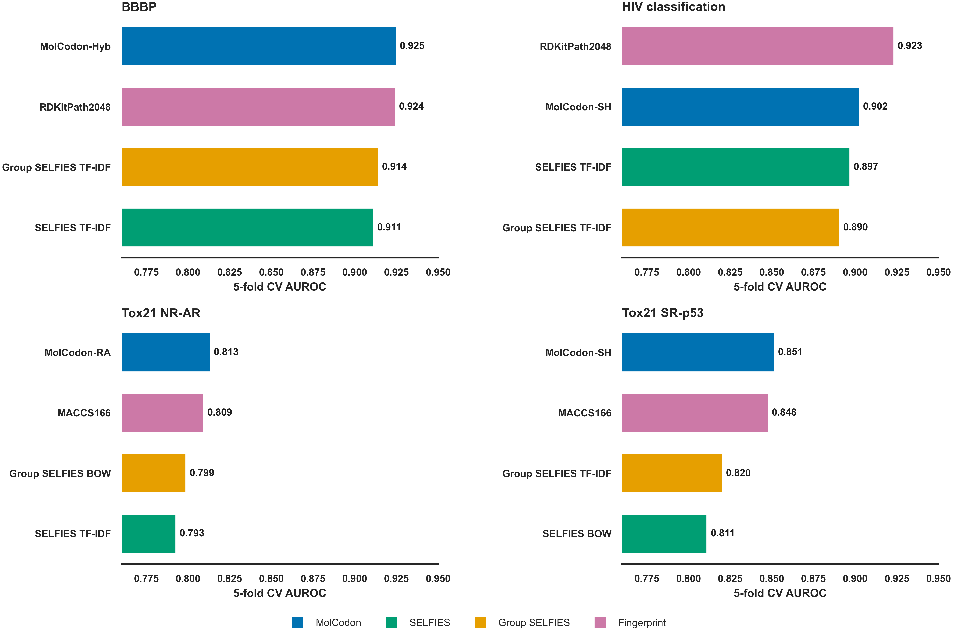
Classification tasks. Best representation within each family on the four binary classification tasks using the best cross-validated learner for each representation. The primary metric is 5-fold CV AUROC.

BBBP showed near-equivalent performance between MolCodon and fingerprint representations, with MolCodon-Hyb giving a slightly higher cross-validated AUROC than RDKit path-2048 (0.925 vs 0.924). Similar small advantages were observed on the toxicity endpoints. MolCodon-RA achieved an AUROC of 0.813 on Tox21 NR-AR, compared with 0.809 for MACCS166, and MolCodon-SH achieved 0.851 on Tox21 SR-p53, compared with 0.848 for MACCS166. Although these margins are modest, they are consistent across the toxicity endpoints and suggest that MolCodon encodes sequence and trace-level features relevant to these classification tasks. For HIV classification, RDKit path-2048 gave the highest AUROC (0.923), while MolCodon-SH remained close at 0.902, indicating that path-based fingerprints are particularly effective for this endpoint but that MolCodon retains competitive performance as a sequence-derived representation. ROC and precision–recall diagnostics for the classification tasks are shown in Figure 6.

**Figure 6:**
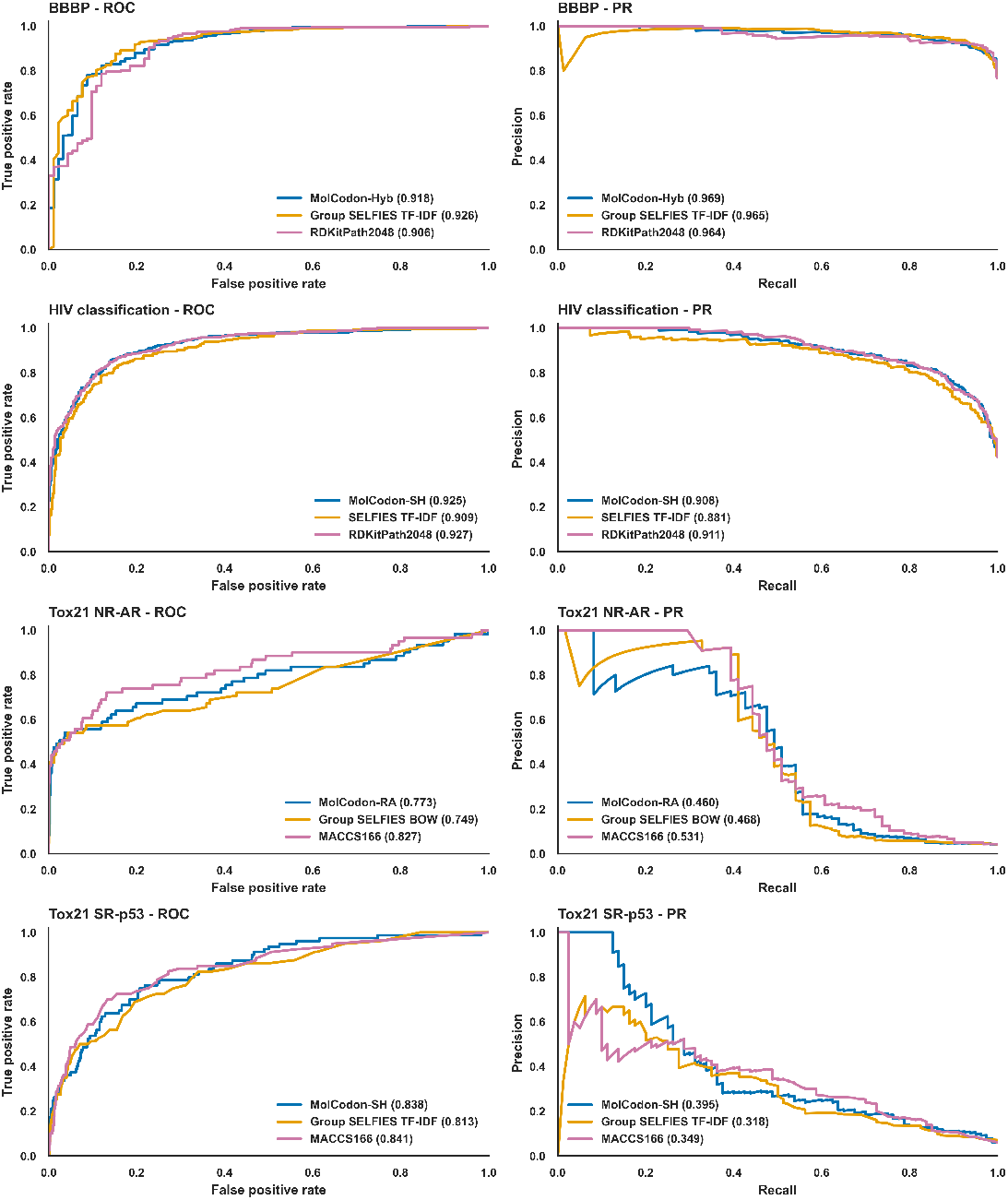
Classification ROC and precision–recall diagnostics. Curves use stored test-set prediction files for the best available MolCodon, SELFIES-family, and fingerprint representation– model pair within each classification task. ROC curves summarize ranking performance, while precision–recall curves emphasize positive-class retrieval in class-imbalanced settings.

The ROC and precision–recall curves provide a complementary view of the classification results and highlight the effect of class imbalance. For BBBP and Tox21 SR-p53, the MolCodon models produced curves that were closely aligned with, and in some regions above, the strongest alternative representations. For HIV classification, the fingerprint model retained an advantage across much of the ROC and precision–recall space, although MolCodon preserved a competitive ranking profile. For Tox21 NR-AR, the precision–recall analysis was especially useful: AUROC values differed only modestly, whereas positive-class retrieval showed clearer separation among representations. (Figure 7)

**Figure 7:**
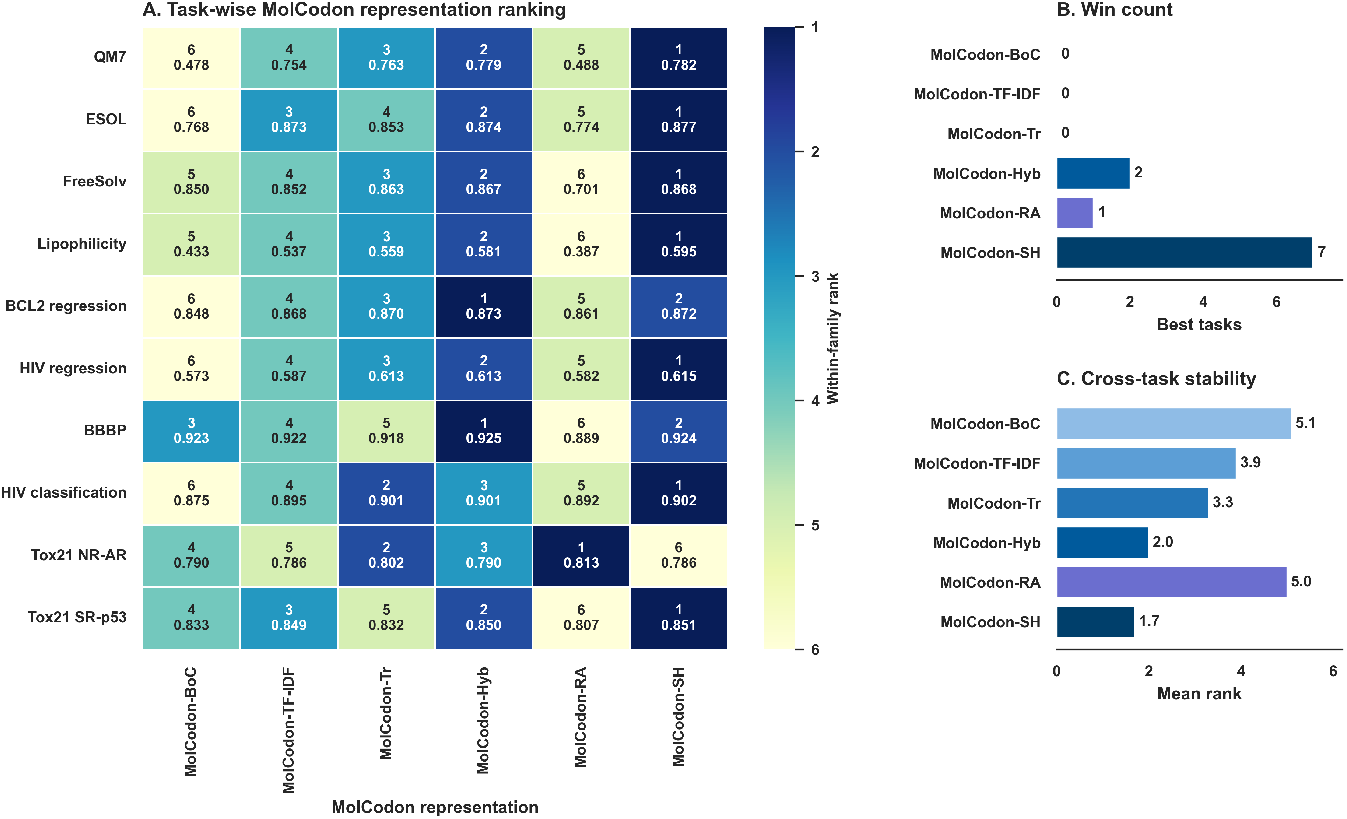
Internal comparison of MolCodon representation views. Panel A shows the within-MolCodon rank for each representation on each endpoint, with the task-primary cross-validation score shown below the rank. Panel B summarizes the number of endpoints won by each representation, and Panel C summarizes mean within-family rank across the ten endpoints. Regression tasks use CV *R*^2^, and classification tasks use CV AUROC.

#### Internal Comparison of MolCodon Representation

The six MolCodon representation showed a clear hierarchy but not a single universally dominant representation (Figure 7). MolCodon-SH was the best MolCodon representation on seven of ten endpoints, including QM7, ESOL, FreeSolv, Lipophilicity, HIV regression, HIV classification, and Tox21 SR-p53. MolCodon-Hyb was the best representation on BCL2 regression and BBBP, while MolCodon-RA was best on Tox21 NR-AR. MolCodon-SH also had the strongest mean within-family rank across endpoints (1.7), followed by MolCodon-Hyb (2.0). This pattern indicates that combining sequence, trace, and role-aware information is usually beneficial, although endpoint-specific MolCodon representation can still be preferable.

The single-channel representations were more endpoint-specific. MolCodon-RA was best on Tox21 NR-AR, suggesting that explicit matched ring and branch contexts can be useful for selected toxicity endpoints. By contrast, MolCodon-BOC and MolCodon-TF–IDF were rarely top-ranked inside the MolCodon family, although they remained informative baselines for isolating the incremental contribution of MolCodon-Tr and MolCodon-RA features. An ablation-style comparison against the MolCodon-BOC baseline is provided in Supplementary Figure S6. That analysis confirms that sequence weighting and trace/hybrid channels account for most gains over the binary bag-of-codons representation, while MolCodon-RA behaves as a more selective endpoint-specific extension.

#### Overview of Family-Level Performance

The family-level summary supports the two main QSAR observations of this benchmark (Figure 8). First, MolCodon outperformed the best SELFIES-family representation on all ten tasks, covering both regression and classification. This comparison isolates the effect of string representation design, because MolCodon, SELFIES, and Group SELFIES are all token-based molecular languages, whereas MolCodon additionally exposes chemical annotations, trace-derived graph events, and role-aware structural context.

**Figure 8:**
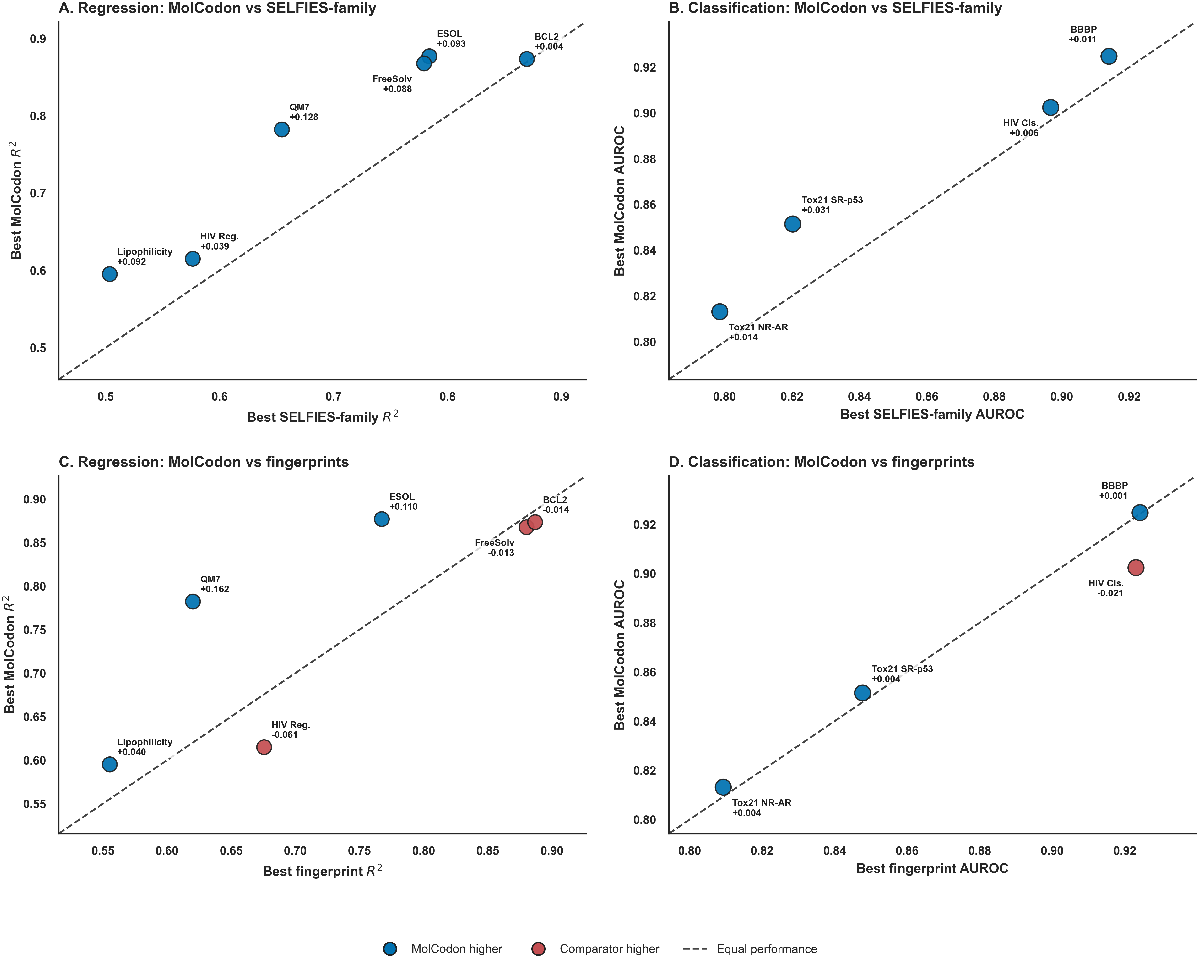
Task-wise family-level comparison of MolCodon against string and fingerprint baselines. Panels A and B compare the best MolCodon representation–model pair with the best SELFIES-family representation–model pair for regression and classification tasks, respectively. Panels C and D compare the best MolCodon representation–model pair with the best fingerprint representation–model pair for the same regression and classification tasks. Regression panels use 5-fold CV *R*^2^, and classification panels use 5-fold CV AUROC. Points above the diagonal indicate higher MolCodon performance. Labels report the task and MolCodon-minus-baseline difference.

Second, the comparison with classical fingerprint baselines was task-dependent rather than uniformly favorable to one representation family. Under the family-best representation–model analysis, MolCodon achieved the best score on six tasks, namely QM7, ESOL, Lipophilicity, BBBP, Tox21 NR-AR, and Tox21 SR-p53. Fingerprints led four tasks, namely FreeSolv, BCL2 regression, HIV regression, and HIV classification. Thus, MolCodon consistently improved over SELFIES-style string baselines, while complementing rather than fully replacing classical fingerprints.

The largest MolCodon advantages over SELFIES-family baselines occurred on QM7 (+0.128 CV *R*^2^), ESOL (+0.093), Lipophilicity (+0.092), and FreeSolv (+0.088). Classification gains over SELFIES-family representations were smaller but remained positive, ranging from +0.006 AUROC on HIV classification to +0.031 AUROC on Tox21 SR-p53. Against the best fingerprint baseline, the largest positive MolCodon effects were observed on QM7 (+0.162 CV *R*^2^ versus RDKit path-2048) and ESOL (+0.110 CV *R*^2^ versus MACCS166). The strongest opposing case was HIV regression, where RDKit path-2048 exceeded MolCodon-SH by 0.061 CV *R*^2^. A chemical-space diagnostic of the random 80:20 partitions is provided in Supplementary Figure S8. The same benchmark was also repeated under Bemis–Murcko scaffold splitting as a secondary robustness analysis, with results provided in Supplementary Figures S2 and S3. The corresponding task-wise delta-bar summaries are provided in Supplementary Figures S4–S7.

### 3.4 PARP1 Virtual Screening

#### Hit Retrieval and Scaffold Diversity

Using olaparib as the reference query, MolCodon BLAST searches of the FDA-approved drug library and the SPECS 330K library returned ranked candidate sets of 50 compounds from each collection. Bemis–Murcko scaffold analysis of the FDA-library hits identified 34 distinct scaffolds, indicating that the search did not simply recover close scaffold analogues of olaparib but also sampled structurally diverse chemotypes. Despite this scaffold diversity, the hits retained substantial correspondence to the query in chemically relevant regions, with median Ring F1 and Pharmacophore F1 scores of 72.3% and 68.9%, respectively. These results suggest that MolCodon BLAST can support scaffold-hopping searches by identifying compounds with different core structures while preserving key ring-topological and pharmacophore features associated with the reference molecule.

#### Physics-based Molecular Simulations

Glide XP docking of the two MolCodon BLAST candidate sets into the PARP1 catalytic site showed distinct score distributions. Compounds from the SPECS library gave more favorable docking scores than the FDA-approved drug candidates, with a median of approximately -7.0 kcal/mol compared with approximately -4.5 to -5.0 kcal/mol for the FDA set. This difference is consistent with the broader and more chemically diverse search space represented by the SPECS library. The ten highest-ranked compounds from each candidate set were selected for MD simulations and are listed in Table 4.

#### Molecular Dynamics and MM/GBSA Binding Free Energies

MM/GBSA binding free energies computed from 20 ns all-atom MD simulations (3 replicas) are presented per compound in Table 4 and as dataset-level distributions in Figure 3.

Docking scores and MM/GBSA binding free energies were initially assessed for the reference PARP inhibitors in order to create a strict comparable baseline. With a docking score of -15.59 kcal/mol and an MM/GBSA energy of -100.67 kcal/mol, olaparib demonstrated the strongest target affinity. Talazoparib and saruparib, two more clinical references, produced MM/GBSA values of -59.13 kcal/mol (docking = -5.30 kcal/mol) and -39.35 kcal/mol (docking = -2.88 kcal/mol), respectively. The newly screened datasets displayed unique MM/GBSA profiles against this established pharmacological spectrum. Among all the screened possibilities, AK-968/40726826 (-74.3 kcal/mol), a molecule with Tanimoto = 0.21 to Olaparib, was the strongest SPECS binder. Scaffold-hopping to a novel chemotype with good binding free energy was successfully demonstrated; it significantly outperformed the affinities of both talazoparib and saruparib. A wider distribution of binding energies (median -51.4 kcal/mol) was obtained using the SPECS–MolCodon method.

FDA-MolCodon displayed the highest MM/GBSA distribution (range -33.7 to -78.5 kcal/mol), which is directly associated with the higher scaffold diversity of MolCodon-retrieved hits. Importantly, Oxatomide (-78.5 ± 4.2 kcal/mol), the top FDA–MolCodon hit, performed better in MM/GBSA than all other screened compounds, placing its binding stability well above talazoparib and saruparib and very close to olaparib’s affinity. Oxatomide, an antihistamine with a piperazine-phthalimide scaffold (Tanimoto = 0.24 to Olaparib), was found because its pharmacophore pattern and ring structure match the Olaparib encoding despite global fingerprint dissimilarity.

These results indicate that MolCodon BLAST can identify binding-competent candidates beyond the scaffold neighborhoods typically prioritized by Tanimoto similarity alone. By incorporating pharmacophore correspondence and ring-topology similarity into the scoring function, MolCodon BLAST retains features that are directly relevant to target engagement while allowing greater variation in the underlying molecular scaffold. This behavior is consistent with the intended use of the method for scaffold-hopping searches, where the goal is not only to recover close analogues but also to identify structurally distinct compounds that preserve key interaction patterns.

The broad MM/GBSA distribution observed for the MolCodon-ranked hits reflects this wider chemical sampling. Rather than concentrating exclusively on compounds with high fingerprint similarity to the reference ligand, the method prioritizes candidates that maintain compatible ring and pharmacophore organization despite lower conventional structural similarity. Notably, individual scaffold-hopping candidates such as Oxatomide and AK-968/40726826 reached favorable binding free-energy estimates, although they would be less likely to be prioritized by a screening strategy based only on fingerprint Tanimoto similarity. These findings support the utility of MolCodon BLAST as a complementary similarity-search framework for identifying chemically diverse, binding-competent candidates.

#### MolCodon BLAST Score Interpretability

Oxatomide provides a useful example of how the MolCodon BLAST scoring scheme can recover scaffold-hopping candidates that have low conventional fingerprint similarity to the query. Although its Tanimoto similarity to olaparib was only 0.24, its decomposed MolCodon scores were high for several chemically relevant components: Ring F1 = 87.5, Branch F1 = 66.7, Attachment F1 = 72.3, Pharmacophore F1 = 91.2, and Overall = 78.0. This score profile indicates that the match was driven primarily by conserved ring topology and pharmacophore organization rather than by close local substructure overlap.

The structural basis for this result is clear from the component-level comparison. Olaparib contains a phthalazinone core, whereas Oxatomide contains a phthalimide core. These scaffolds are not close analogues in a conventional fingerprint sense, but they preserve a related ring-topological arrangement and a similar pharmacophore pattern, including an NH hydrogen-bond donor and a ring oxygen hydrogen-bond acceptor. MolCodon BLAST assigns explicit credit to these shared features through its ring and pharmacophore terms, while still accounting for differences in branch and attachment architecture.

By contrast, circular fingerprint Tanimoto similarity is dominated by overlap in local atom-centered environments. As a result, it may assign a low similarity score to molecules that differ in local fragment composition even when they preserve higher-level ring organization and interaction-relevant pharmacophore features. The advantage of the MolCodon BLAST output in this case is therefore not only the ranking itself, but also its interpretability. The overall similarity score can be decomposed into ring, branch, attachment, and pharmacophore contributions, providing a direct structural rationale for why a low-Tanimoto compound was selected.

Figure 9 shows the MolCodon BLAST comparison between Olaparib and Oxatomide, the top-ranked FDA-library MolCodon hit by MM/GBSA analysis (-78.5 kcal/mol). Although Oxatomide showed low conventional fingerprint similarity to Olaparib (Tanimoto = 0.23), MolCodon BLAST assigned an overall score of 56.29. This score was driven mainly by strong pharmacophore conservation, with a Pharmacophore F1 of 87.50, corresponding to 7 of 8 pharmacophore features matched in comparable structural contexts. The component-level breakdown clarifies the basis of the match. Three of the five ring systems were matched, giving a Ring F1 of 66.67. These include the piperazine ring and two carbocyclic aromatic systems that are retained across the phthalazinone-containing scaffold of Olaparib and the benzimidazolone/diphenylmethyl framework of Oxatomide. The remaining differences arise from ring systems that are not directly conserved, including the cyclopropane carboxamide region of Olaparib and the fused benzimidazole-containing region of Oxatomide. These unmatched elements account for the lower Branch F1 and Attachment F1 values of 54.55 and 36.36, respectively.

**Figure 9:**
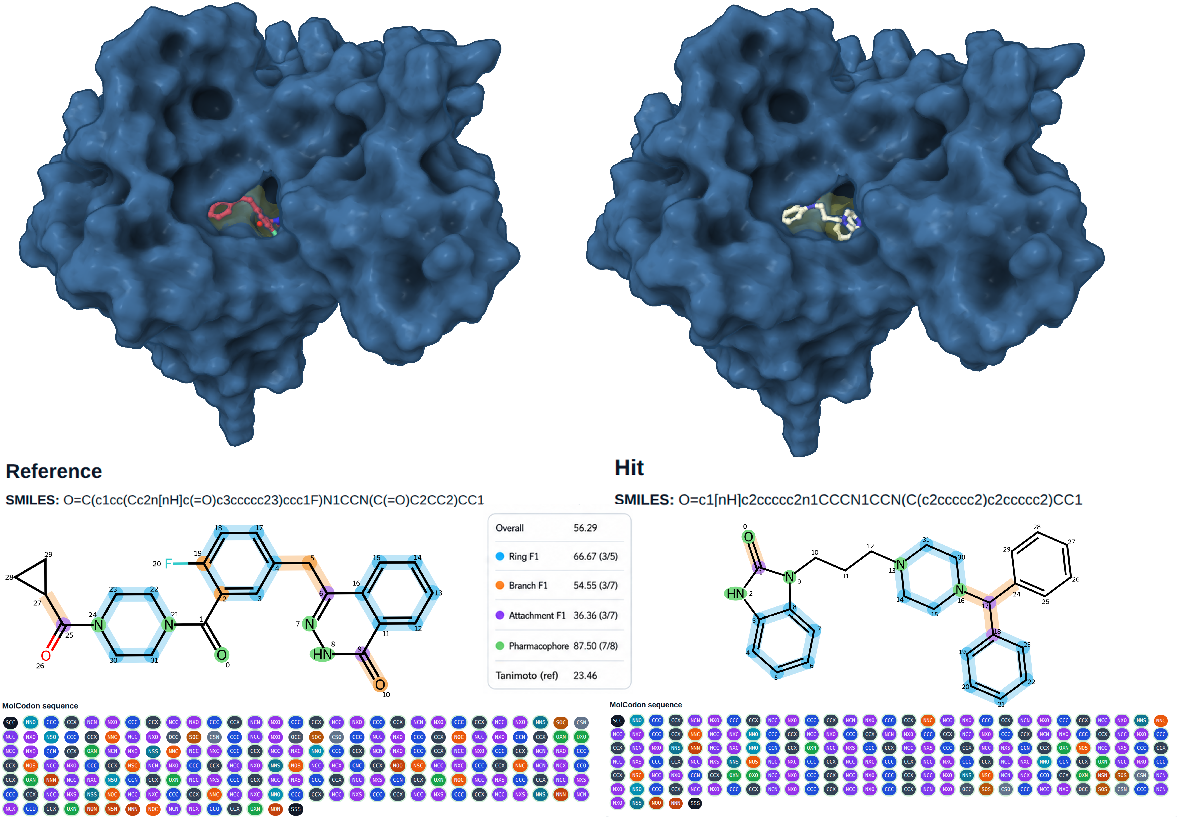
MolCodon BLAST scaffold-hopping case study: Olaparib (reference) versus Oxatomide (FDA–MolCodon hit). Upper panels: Glide XP docking poses of Olaparib (left, red/green sticks) and Oxatomide (right, white sticks) in the PARP1 NAD^+^-binding site (PDB: 7KK4; protein surface in blue). Both ligands occupy the same catalytic cleft despite their structural dissimilarity (Tanimoto = 0.23). Lower panels: MolCodon component-match visualizations generated by molcodon_viz.py. Matched ring systems are highlighted in blue on the 2D structures; matched branches in orange; matched pharmacophore features in green. The full codon sequence for each molecule is shown beneath the structure, with codon tokens colour-coded by family. Score card (centre): Overall = 56.29; Ring F1 = 66.67 (3/5 rings matched); Branch F1 = 54.55 (3/7); Attachment F1 = 36.36 (3/7); Pharmacophore F1 = 87.50 (7/8 features matched); Tanimoto (ref) = 23.46. The near-complete pharmacophore match, driven by conserved HBA/HBD pattern in matching structural contexts, is the primary signal that caused MolCodon BLAST to rank Oxatomide highly despite its low fingerprint similarity.

The docking poses shown in the upper panels of Figure 9 further support this interpretation, with Oxatomide occupying the PARP1 *NAD*^+^-binding cleft in a pose comparable to that of Olaparib. This example illustrates the scaffold-hopping behavior of MolCodon BLAST. Rather than relying solely on local circular-fragment overlap, the method can prioritize compounds that preserve pharmacophore arrangement and ring-topological correspondence despite substantial scaffold divergence. In this case, MolCodon BLAST identified a clinically approved antihistamine as a structurally distinct PARP1-binding candidate. A corresponding analysis for the top SPECS-library MolCodon hit, AK/968/40726826, is provided in Figure S9.

## 4. Discussion

The internal comparison of MolCodon feature views reinforces this interpretation. Representations that combined sequence n-grams with trace and role-aware information generally performed better than simple bag-of-codon features. This suggests that the most useful signal is not merely the presence of individual codons, but the combination of codon order, structural role, ring and branch context, and graph-derived annotations. In this respect, MolCodon offers a natural bridge between molecular language models and graph descriptors. It keeps the computational convenience of sparse sequence features while retaining a direct connection to structural events in the molecular graph.

The PARP1 virtual screening case study illustrates a complementary use of MolCodon: interpretable scaffold hopping. Conventional fingerprint similarity is highly effective for identifying close analogues, but it can under-rank molecules that preserve pharmacophore arrangement or ring-topological correspondence while differing in local circular fragments. MolCodon BLAST was designed for this setting. By decomposing similarity into ring, branch, attachment, and pharmacophore components, it can identify molecules that are not close fingerprint analogues but still preserve chemically relevant features of the query. The olaparib screening experiment demonstrates this behavior. The FDA-library hits contained substantial scaffold diversity, and individual low-Tanimoto candidates were nevertheless predicted to occupy the PARP1 *NAD*^+^-binding cleft with favorable computed binding energetics.

Oxatomide is the clearest example. Despite low fingerprint similarity to olaparib, MolCodon BLAST prioritized this compound because it retained a strong pharmacophore match and meaningful ring-topological correspondence. The component-level score decomposition explains the result in chemical terms: the overall match is driven by conserved hydrogen-bonding patterns and ring architecture rather than by local fragment overlap alone. This type of explanation is difficult to obtain from a single Tanimoto coefficient. The identification of AK-968/40726826 from the SPECS library provides a second example in which MolCodon retrieved a structurally distinct candidate with favorable computed binding free energy. These results support MolCodon BLAST as a practical scaffold-hopping tool, particularly when the goal is to explore chemically diverse analogues that preserve key interaction patterns.

### Limitations

The present study also defines clear boundaries for the current implementation and points to practical opportunities for extension. The PARP1 docking, MD, and MM/GBSA analyses provide structure-based support for the MolCodon BLAST hits, but these results should be interpreted as computational prioritization rather than direct evidence of biological activity. Within this context, the simulations were used to assess whether MolCodon-retrieved scaffold-hopping candidates could adopt stable and energetically favorable binding modes in the PARP1 catalytic pocket. Longer simulations and experimental assays would be valuable next steps for validating the most promising candidates. The current MolCodon alphabet was intentionally designed around drug-like organic chemistry and therefore does not yet cover all elements, isotope labels, or highly specialized chemical classes. Similarly, the present ring-encoding strategy performs well across large screening libraries but has defined boundaries for highly fused or bridged polycyclic systems. These limitations are largely scope-related and can be addressed by extending the codon vocabulary and ring-reference logic. Finally, although the QSAR benchmark spans diverse physicochemical, toxicity, permeability, and bioactivity endpoints, additional prospective datasets and target-specific evaluations will further clarify where MolCodon provides the greatest predictive and interpretive benefit.

### Future Directions

These limitations also define clear future directions. Expanding the codon vocabulary to support additional elements, isotopes, organometallic motifs, and more complex ring systems would broaden chemical coverage while preserving the fixed-width design. The explicit trace structure of MolCodon also makes it well suited for transformer-based molecular language models, where attention weights could be related back to codon families, ring systems, branch contexts, or pharmacophore annotations. In generative modeling, the same codon grammar could be used to impose chemically meaningful constraints during sequence generation. For similarity search, target-specific or activity-learned component weights could adapt MolCodon BLAST to particular protein families or phenotypic screens. Finally, integration with 3D pharmacophore matching, docking rescoring, or active-learning workflows could further improve enrichment while preserving the interpretability of the retrieval process.

## 5. Conclusion

MolCodon introduces a fixed-width, chemically annotated molecular language that creates a direct interface between molecular graph representation and biological sequence analysis. Its novelty lies in representing small molecules as deterministic three-character codon sequences over a compact symbolic alphabet, while assigning each codon family a defined chemical function. Atoms, bonds, rings, branches, fused-ring references, formal charge, stereochemistry, bond mobility, and pharmacophore features are therefore encoded as explicit and interpretable sequence elements rather than being embedded in traversal syntax, hashed into fingerprint bits, or compressed into latent vectors. This design preserves a direct connection between the sequence and the underlying molecular graph, while its ring-contiguous organization keeps cyclic scaffolds locally organized in the encoded string. As a result, MolCodon provides not only a molecular notation, but a structurally decomposable language in which sequence tokens can be traced back to chemically meaningful graph events. This combination of deterministic encoding, high-fidelity reconstruction, explicit chemical annotation, and compatibility with sequence-based analysis distinguishes MolCodon from existing molecular representations and establishes a new framework for interpretable similarity search, QSAR feature generation, scaffold exploration, and bioinformatics-inspired analysis of small-molecule chemical space.

## Data and Software Availability

Open-source software for the MolCodon encoder, decoder, BLAST similarity engine, trace annotator, and visualization scripts can be found at https://github.com/DurdagiLab/MolCodon. PDB structure 7KK4: https://www.rcsb.org/.

## Acknowledgements

This study was funded by the Scientific Research Projects Commission of Bahçeşehir University, project numbers BAP.2024.01.42 and BAP.2025-01.52. This study was also funded by Uğur-HİTMER Research Fund.

## Conflict of Interest

The authors declare no competing financial interests.

---

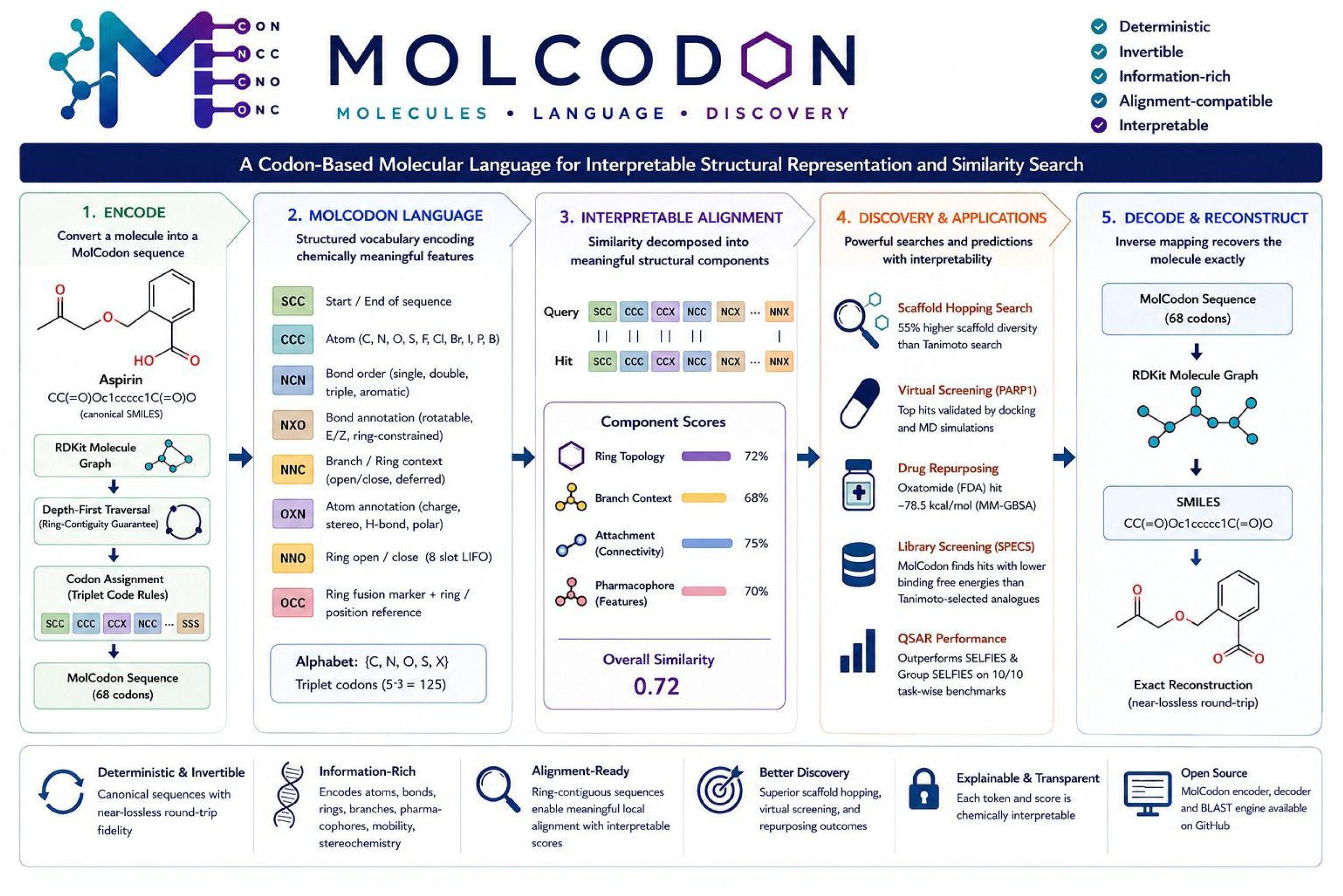

## Notes

### Competing Interest Statement

The authors have declared no competing interest.

