## Supporting Material for "MolCodon: A Codon-Based Molecular Language for Interpretable Structural Representation and Similarity Search"

[*]Corresponding author: Serdar Durdağı

May 20, 2026

#### S1 MolCodon Codon Dictionary

Table S1: Complete MolCodon codon dictionary. Each row gives the three-letter codon, its structural meaning, and its functional category. Ring and branch codons come in matched open/close pairs that share a label slot; e.g. NNO opens ring slot 0 and NNS closes it. The encoder maintains an 8-slot LIFO for ring labels and a 4-slot rolling counter for branch labels, which is why the dictionary contains exactly 16 ring codons and 8 branch codons.

| # | Codon | Meaning | Category |
|---|---|---|---|
| 1 | SCC | Start of sequence – first codon of every encoded molecule | Special |
| 2 | SSS | End of sequence – last codon of every encoded molecule | Special |
| 3 | OCC | Ring fusion marker – followed by a ring-reference + position-reference codon pair | Special |
| 4 | CCC | Carbon (C) | Atom |
| 5 | CCN | Nitrogen (N) | Atom |
| 6 | CCO | Oxygen (O) | Atom |
| 7 | CCS | Sulfur (S) | Atom |
| 8 | CNC | Fluorine (F) | Atom |
| 9 | CNN | Chlorine (Cl) | Atom |
| 10 | CNO | Bromine (Br) | Atom |
| 11 | CNS | Iodine (I) | Atom |
| 12 | COC | Phosphorus (P) | Atom |
| 13 | CON | Boron (B) | Atom |
| 14 | NCC | Single bond (– ) | Bond order |
| 15 | NCN | Double bond (=) | Bond order |
| 16 | NCO | Triple bond (≡) | Bond order |
| 17 | NCS | Aromatic bond | Bond order |
| 18 | CCX | Formal charge = 0 (neutral) | Atom annotation (charge) |
| 19 | CXN | Formal charge = +1 | Atom annotation (charge) |
| 20 | CXS | Formal charge $\geq +2$ | Atom annotation (charge) |
| 21 | CXO | Formal charge = −1 | Atom annotation (charge) |
| 22 | CXX | Formal charge $\leq -2$ | Atom annotation (charge) |
| 23 | SXN | CIP stereo = R | Atom annotation (stereo) |
| 24 | SXO | CIP stereo = S | Atom annotation (stereo) |
| 25 | OXN | Hydrogen-bond acceptor, emitted for every N and O atom | Atom annotation (pharmacophore) |
| 26 | OXO | Polar hydrogen, emitted once per H bound to N or O | Atom annotation (pharmacophore) |
| 27 | NXC | Rotatable single bond | Bond annotation (mobility) |

Continued on next page

Table S1 – continued from previous page

| # | Codon | Meaning | Category |
|---|---|---|---|
| 28 | NCX | Non-rotatable terminal single, double, or triple bond | Bond annotation (mobility) |
| 29 | NXS | Ring-constrained non-aromatic ring bond | Bond annotation (mobility) |
| 30 | NXO | Aromatic-locked bond in aromatic ring | Bond annotation (mobility) |
| 31 | SOX | Double-bond stereo = E (trans) | Bond annotation (stereo) |
| 32 | SNX | Double-bond stereo = Z (cis) | Bond annotation (stereo) |
| 33 | NNC | Branch open (label 0) | Branch |
| 34 | NNN | Branch close (label 0) | Branch |
| 35 | NOC | Branch open (label 1) | Branch |
| 36 | NON | Branch close (label 1) | Branch |
| 37 | NOS | Branch open (label 2) | Branch |
| 38 | NOO | Branch close (label 2) | Branch |
| 39 | NSC | Branch open (label 3) | Branch |
| 40 | NSN | Branch close (label 3) | Branch |
| 41 | NNO | Ring open (slot 0) | Ring |
| 42 | NNS | Ring close (slot 0) | Ring |
| 43 | NSO | Ring open (slot 1) | Ring |
| 44 | NSS | Ring close (slot 1) | Ring |
| 45 | OSO | Ring open (slot 2) | Ring |
| 46 | OSS | Ring close (slot 2) | Ring |
| 47 | OCN | Ring open (slot 3) | Ring |
| 48 | ONO | Ring close (slot 3) | Ring |
| 49 | OCO | Ring open (slot 4) | Ring |
| 50 | ONS | Ring close (slot 4) | Ring |
| 51 | OCS | Ring open (slot 5) | Ring |
| 52 | OSC | Ring close (slot 5) | Ring |
| 53 | ONC | Ring open (slot 6) | Ring |
| 54 | OSN | Ring close (slot 6) | Ring |
| 55 | ONN | Ring open (slot 7) | Ring |
| 56 | SCN | Ring close (slot 7) | Ring |
| 57 | SOC | Ring reference – completed ring index 0 | Reference |
| 58 | SON | Ring reference – completed ring index 1 | Reference |
| 59 | SOO | Ring reference – completed ring index 2 | Reference |
| 60 | SOS | Ring reference – completed ring index 3 | Reference |
| 61 | SSC | Ring reference – completed ring index 4 | Reference |
| 62 | SSN | Ring reference – completed ring index 5 | Reference |
| 63 | SSO | Ring reference – completed ring index 6 | Reference |
| 64 | COO | Position reference – atom index 0 within ring | Reference |
| 65 | COS | Position reference – atom index 1 within ring | Reference |
| 66 | CSC | Position reference – atom index 2 within ring | Reference |
| 67 | CSN | Position reference – atom index 3 within ring | Reference |
| 68 | CSO | Position reference – atom index 4 within ring | Reference |
| 69 | CSS | Position reference – atom index 5 within ring | Reference |
| 70 | OOC | Position reference – atom index 6 within ring | Reference |

Continued on next page

Table S1 – continued from previous page

| # | Codon | Meaning | Category |
|---|---|---|---|
| 71 | OON | Position reference – atom index 7 within ring | Reference |
| 72 | OOO | Position reference – atom index 8 within ring | Reference |
| 73 | OOS | Position reference – atom index 9 within ring | Reference |
| 74 | SCO | Position reference – atom index 10 within ring | Reference |
| 75 | SCS | Position reference – atom index 11 within ring | Reference |
| 76 | SNC | Position reference – atom index 12 within ring | Reference |
| 77 | SNN | Position reference – atom index 13 within ring | Reference |
| 78 | SNO | Position reference – atom index 14 within ring | Reference |
| 79 | SNS | Position reference – atom index 15 within ring | Reference |

#### S2 Per-Endpoint Notes

**MoleculeNet QM7.** Atomization energies (kcal/mol) on small organic molecules. After parsing, two structures fail SMILES validation. Two more fail MOLCODON encodability. Final benchmark size $n = 6830$.

**MoleculeNet ESOL.** Aqueous solubility (logS) for 1128 small molecules. Eleven duplicates collapse during canonicalization. Final $n = 1117$, full MOLCODON coverage.

**MoleculeNet FreeSolv.** Hydration free energy (kcal/mol) for 642 molecules. No duplicates and full MOLCODON coverage. It is the smallest dataset in the benchmark.

**MoleculeNet Lipophilicity.** octanol–water distribution coefficient $\log D_{7.4}$ for 4200 molecules. Four molecules are filtered by MOLCODON encodability (transition metals).

**BCL2 Regression.** Curated from ChEMBL with single-target activity for BCL2 inhibition. The cleaned table is in `ChEMBL_BCL2_1067.csv`, and the regression target is $\log(IC_{50}/nM)$. Out of 1068 raw rows, one fails parsing, fourteen are MOLCODON-incompatible (mostly metallic ligands), final $n = 1053$.

**HIV Regression.** Source CSV is `All_HIV_Cleaned_Regression.csv` with a NORMPIC50 column. Two SMILES fail parsing, and twenty-five are not MOLCODON-encodable. final $n = 4208$.

**HIV Classification.** Standard MoleculeNet HIV classification labels. Final $n = 4823$ after deduplication and encodability filtering.

**MoleculeNet BBBP.** Blood–brain-barrier permeability binary labels. Eighty-five canonical-SMILES collisions are merged. Eight molecules fail encodability. Final $n = 1957$.

**Tox21 NR-AR.** Nuclear-receptor androgen-receptor activity. Tox21 is a multitask panel, and we evaluate this single endpoint to sample nuclear-receptor toxicity behaviour. After SMILES parsing, replicate consensus, and MOLCODON encodability filter, $n = 7077$.

**Tox21 SR-p53.** Stress-response p53 activity. Same processing as NR-AR, final $n = 6608$.

The aggregate curation and random-split statistics are reproduced as Table S2. Numbers match those in the main paper.

Table S2: Curation and random-split statistics.

| Task | Raw | Clean | Bench | Train | Test | Cov. |
|---|---|---|---|---|---|---|
| MoleculeNet QM7 | 6834 | 6832 | 6830 | 5464 | 1366 | 1.000 |
| MoleculeNet ESOL | 1128 | 1117 | 1117 | 893 | 224 | 1.000 |
| MoleculeNet FreeSolv | 642 | 642 | 642 | 513 | 129 | 1.000 |
| MoleculeNet Lipophilicity | 4200 | 4200 | 4196 | 3356 | 840 | 0.999 |
| BCL2 Regression | 1068 | 1067 | 1053 | 842 | 211 | 0.987 |
| HIV Regression | 4235 | 4233 | 4208 | 3366 | 842 | 0.994 |
| MoleculeNet BBBP | 2050 | 1965 | 1957 | 1565 | 392 | 0.996 |
| HIV Classification | 4855 | 4852 | 4823 | 3858 | 965 | 0.994 |
| Tox21 NR-AR | 7831 | 7258 | 7077 | 5661 | 1416 | 0.975 |
| Tox21 SR-p53 | 7831 | 6767 | 6608 | 5286 | 1322 | 0.977 |

#### S3 Hyperparameter Grids

Each task–representation pair runs 5-fold cross-validation over the grids in Table S3. Linear models are pre-scaled by `MaxAbsScaler`. Tree models are fit directly on sparse inputs.

Table S3: Hyperparameter grids per candidate model.

| Task type | Model | Grid |
|---|---|---|
| regression | ridge | `alpha` $\in \{0.1, 1.0, 10.0\}$ |
| | linear_svr | `C` $\in \{0.5, 2.0\}$, `epsilon` $\in \{0.01, 0.1\}$ |
| | sgd_svm | `alpha` $\in \{1e{-4}, 1e{-3}\}$ |
| | rbf_svm | `C`=1.0, `epsilon`=0.1, `gamma`=scale |
| | rand. forest | `n_estimators`=400, `max_features` $\in$ {sqrt, 0.5} |
| | lgbm | `n_estimators` $\in \{500\}$, `learning_rate` $\in \{0.05, 0.1\}$, `num_leaves` $\in \{31, 63\}$ |
| | xgboost | `n_estimators` $\in \{500\}$, `learning_rate` $\in \{0.05, 0.1\}$, `max_depth` $\in \{6, 8\}$ |
| classification | logreg | `C` $\in \{0.5, 2.0\}$, `class_weight` $\in$ {none, balanced} |
| | linear_svm | `C` $\in \{0.5, 2.0\}$, `class_weight` $\in$ {none, balanced} |
| | sgd_log | `alpha` $\in \{1e{-4}, 1e{-3}\}$ |
| | sgd_svm | `alpha` $\in \{1e{-4}, 1e{-3}\}$, `class_weight` $\in$ {none, balanced} |
| | rbf_svm | `C`=1.0, `class_weight`=balanced, `gamma`=scale |
| | rand. forest | `n_estimators`=400, `class_weight` $\in$ {none, balanced} |
| | lgbm | `n_estimators` $\in \{500\}$, `learning_rate` $\in \{0.05, 0.1\}$, `num_leaves` $\in \{31, 63\}$ |
| | xgboost | `n_estimators` $\in \{500\}$, `learning_rate` $\in \{0.05, 0.1\}$, `max_depth` $\in \{6, 8\}$ |

#### S4 Model Selection

Borda scoring was computed across the 140 task–representation scenarios using the task-primary cross-validation metric. Regression scenarios were ranked by CV $R^2$, and classification scenarios were ranked by CV AUROC. The matched comparison was restricted to model families available for both regression and classification. XGBoost obtained the highest aggregate Borda score and the three ensemble learners formed the most stable model tier (Table S4). The corresponding diagnostic panel is shown in Figure S1.

Table S4: Model ranking by Borda aggregation.

| Model | Borda | Mean rank | Median rank | First place | Top-2 | Scenarios |
|---|---|---|---|---|---|---|
| XGBoost | 680 | 2.14 | 2.0 | 46 | 94 | 140 |
| LightGBM | 676 | 2.17 | 2.0 | 48 | 91 | 140 |
| Random forest | 630 | 2.50 | 2.0 | 40 | 72 | 140 |
| RBF-SVM | 388 | 4.23 | 4.0 | 3 | 14 | 140 |
| Linear SVM | 297 | 4.88 | 5.0 | 2 | 7 | 140 |
| SGD-SVM | 269 | 5.08 | 5.0 | 1 | 2 | 140 |

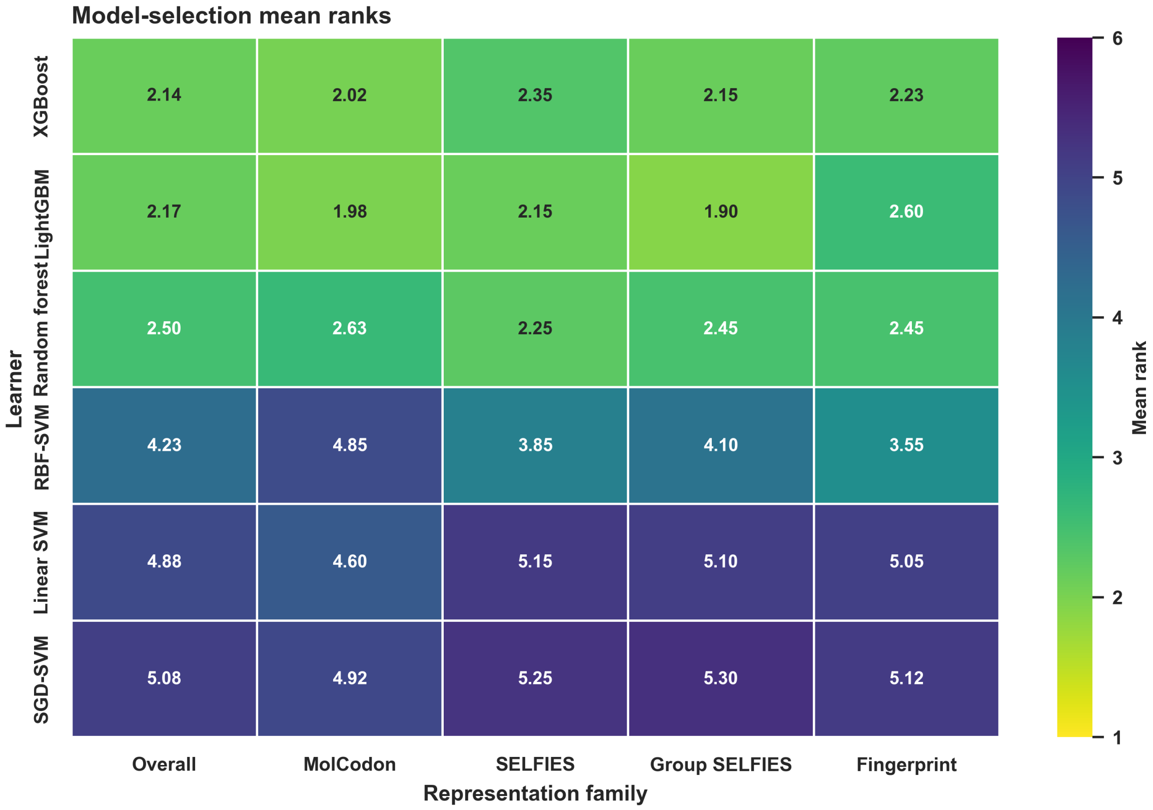


Figure S1: Model-selection diagnostics. Candidate learners were ranked using 5-fold CV $R^2$ for regression tasks and 5-fold CV AUROC for classification tasks. The heatmap reports mean rank overall and by representation family. Overall denotes the mean rank across all 140 task–representation scenarios used for Borda aggregation. Lower rank indicates stronger robustness.

#### S5 Scaffold-Split Robustness Analysis

The main manuscript reports the random 80:20 split as the primary QSAR benchmark. To assess whether the representation-level patterns were overly dependent on random train–test assignment, the full benchmark was repeated with Bemis–Murcko scaffold-based 80:20 splits. In these runs, scaffold groups were assigned to either train or test, and the final scaffold summary contained no shared train/test scaffolds for any endpoint. The same curation, representation, model-selection, and evaluation workflow used for the random-split analysis was then applied to the scaffold-split partitions.

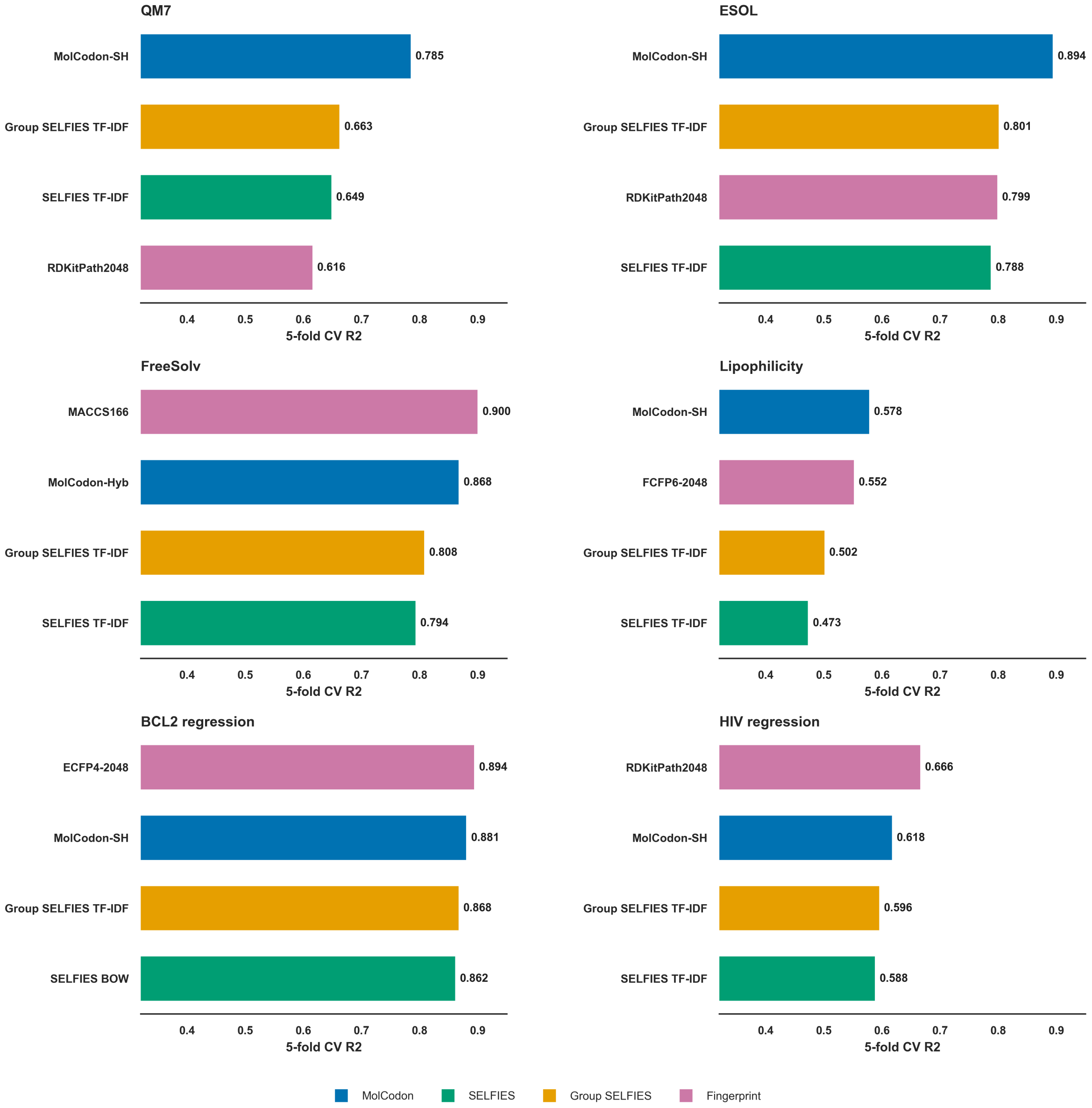


Figure S2: Scaffold-split family-best comparison for the six regression endpoints. Bars show the best representation within each family under the scaffold-split benchmark. Scores are 5-fold CV $R^2$ values computed inside the scaffold-based training partition.

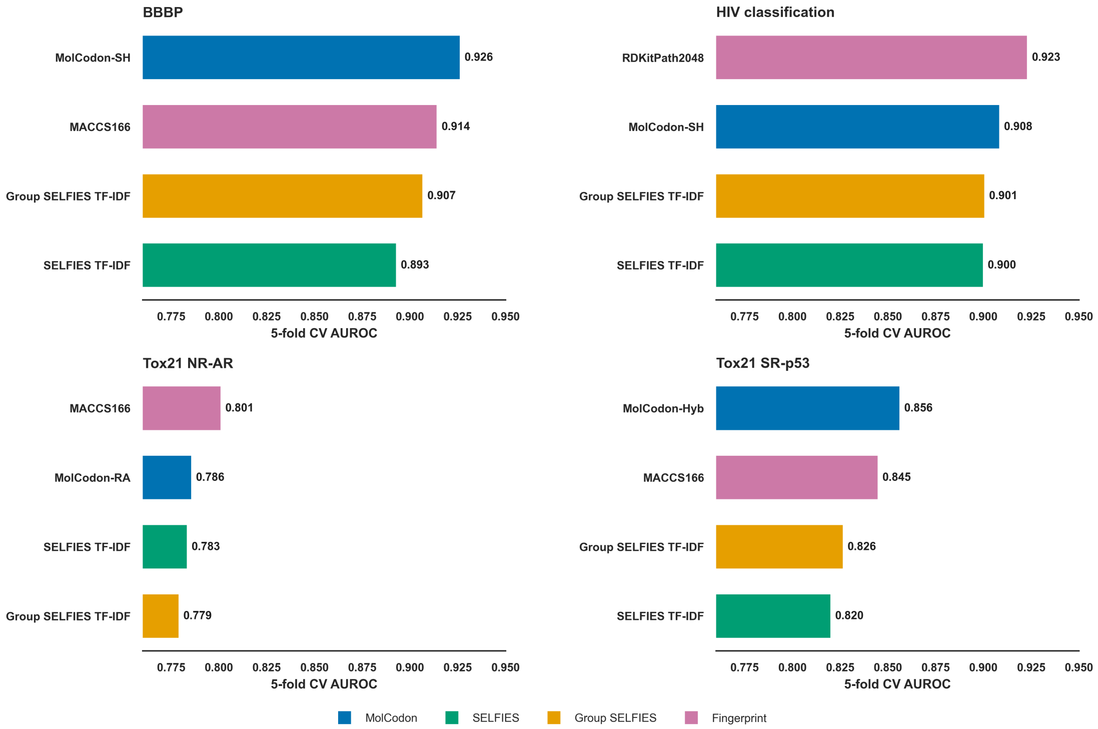


Figure S3: Scaffold-split family-best comparison for the four classification endpoints. Bars show the best representation within each family under the scaffold-split benchmark. Scores are 5-fold CV AUROC values computed inside the scaffold-based training partition.

#### S6 Task-Wise Delta Summary

The main manuscript presents the MolCodon versus best non-MolCodon comparison as a paired scatter plot. Figure S4 provides the same comparison as signed task-wise differences for readers who prefer the direct delta representation.

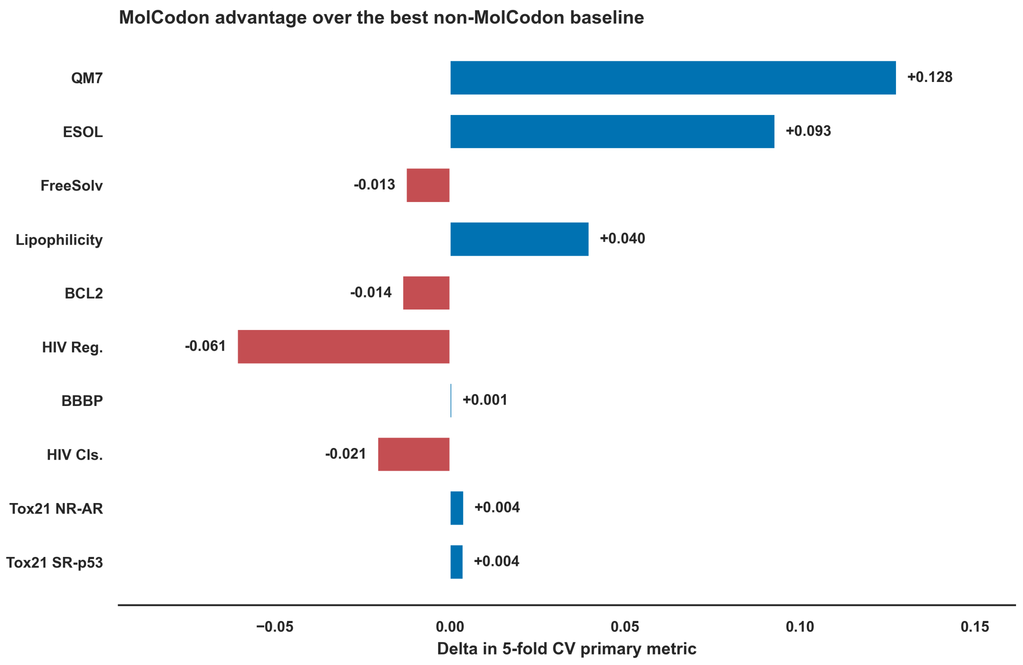


Figure S4: Task-wise delta between the best MolCodon representation and the best non-MolCodon baseline. Positive values indicate a MolCodon advantage. Regression tasks use 5-fold CV $R^2$, and classification tasks use 5-fold CV AUROC.

### S7 MolCodon Versus SELFIES-Family Delta

The main manuscript summarizes the consistent MolCodon advantage over SELFIES and Group SELFIES in text. Figure S5 provides the corresponding task-wise delta panel.

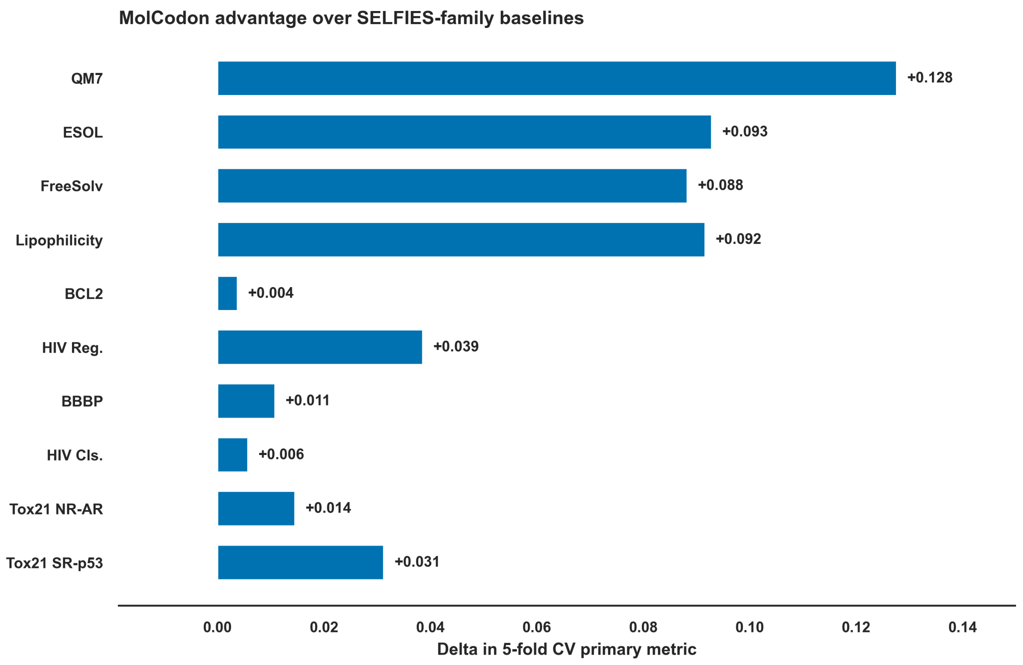


Figure S5: Task-wise delta between the best MolCodon representation and the best SELFIES or Group SELFIES representation. Positive values indicate a MolCodon advantage. Regression tasks use 5-fold CV $R^2$, and classification tasks use 5-fold CV AUROC.

#### S8 MolCodon Representation Ablation

Figure S6 reports the endpoint-wise change in performance relative to MolCodon-BOW. This panel supports the main manuscript's internal MolCodon representation comparison by showing where sequence weighting, trace channels, role-aware channels, and hybrid representations add performance over the binary bag-of-codons baseline.

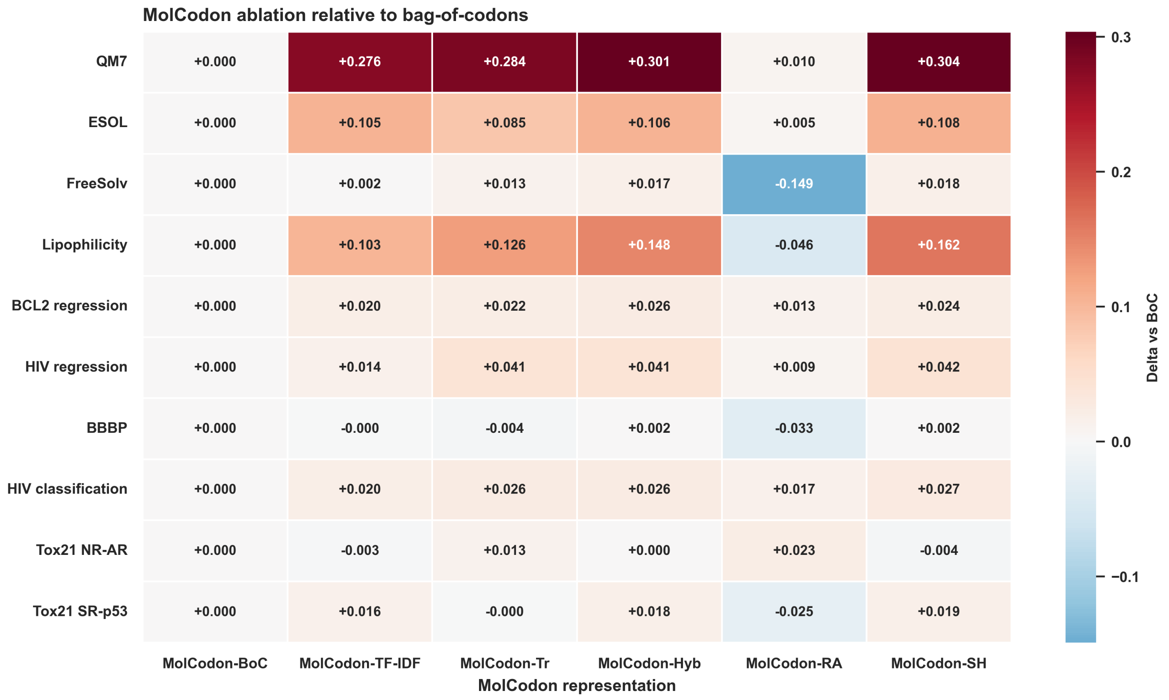


Figure S6: MolCodon representation ablation relative to bag-of-codons. Cells show the change in the task-primary CV metric relative to MolCodon-BOW, followed by the absolute CV score. Regression tasks use 5-fold CV $R^2$, and classification tasks use 5-fold CV AUROC.

#### S9 MolCodon Versus Fingerprint Delta

The main manuscript summarizes the endpoint-dependent comparison between MolCodon and classical fingerprint baselines in text. Figure S7 provides the corresponding task-wise signed deltas.

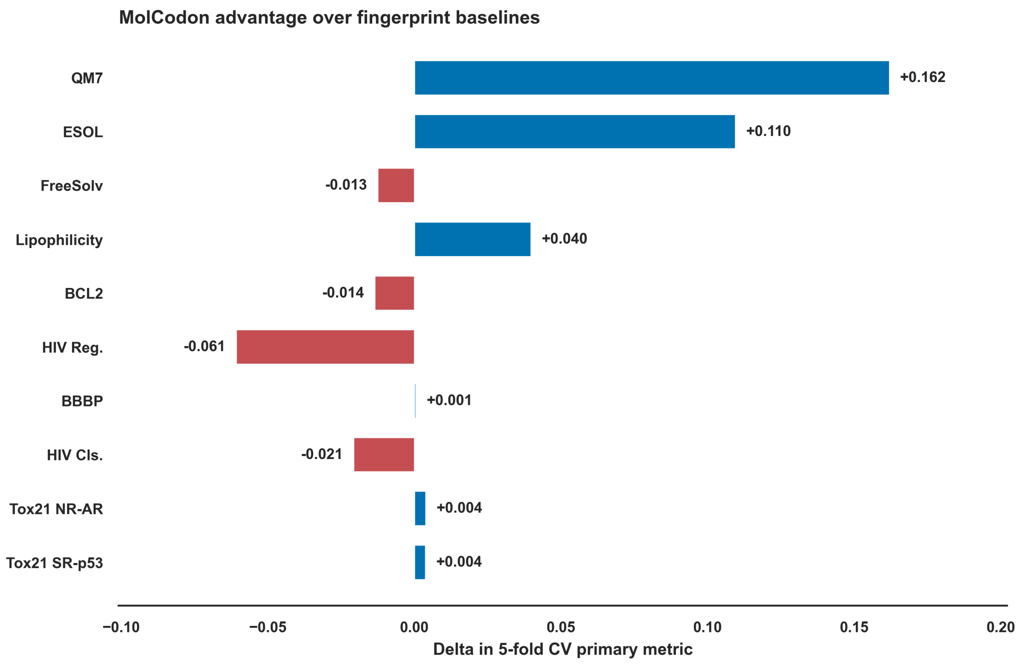


Figure S7: Task-wise delta between the best MolCodon representation and the best fingerprint representation. Positive values indicate a MolCodon advantage. Regression tasks use 5-fold CV $R^2$, and classification tasks use 5-fold CV AUROC.

#### S10 Chemical-Space Coverage of Random Splits

Figure S8 reports the PCA diagnostic used to check whether the random 80:20 partitions preserved broad chemical-space coverage. ECFP4 fingerprints from sampled training and test molecules were embedded by PCA and visualized separately for regression and classification tasks.

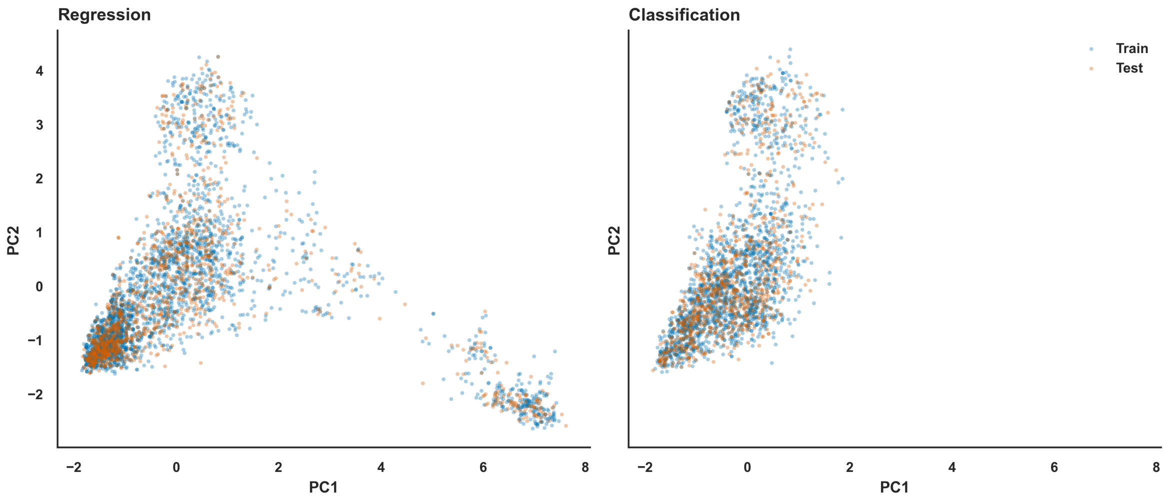


Figure S8: Chemical-space representativeness of random train–test partitions. Molecules were embedded using ECFP4 fingerprints followed by PCA. Circles denote training molecules and triangles denote test molecules. Colors denote benchmark task.

### S11 MolCodon Similarity

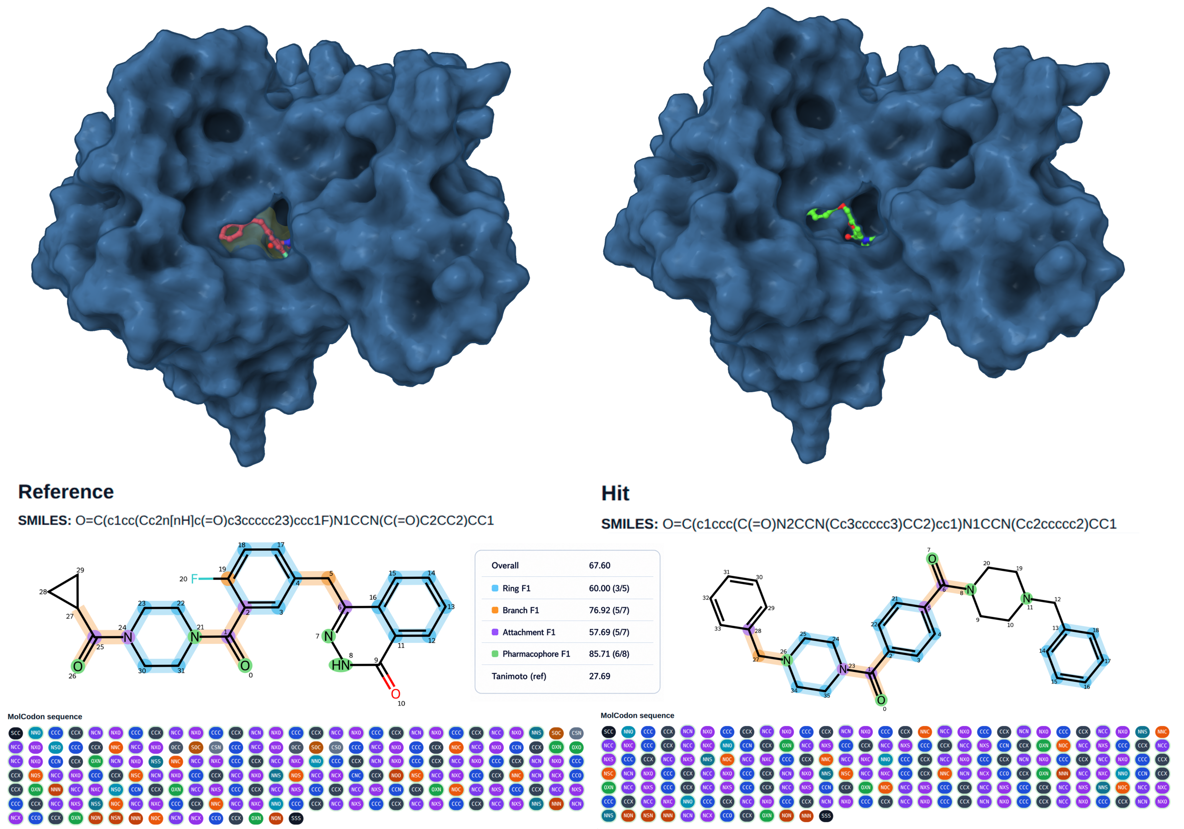


Figure S9: MolCodon BLAST scaffold-hopping case study: Olaparib (reference) versus AK/968/40726826 (SPECS–MolCodon hit). Upper panels: Glide XP docking poses of Olaparib (left) and AK-968/40726826 (right) in the PARP1 NAD$^+$-binding site (PDB: 7KK4; protein surface in blue). Both ligands occupy the same catalytic cleft (MM/GBSA $= -74.3 \pm 5.8$ kcal/mol for AK/968/40726826, 3-replica average over 20 ns). Lower panels: MolCodon matched ring systems are highlighted in blue on the 2D structures; matched branches in orange; matched pharmacophore features in green. The full codon sequence for each molecule is shown beneath the structure, with tokens colour-coded by family.

### S12 Complete Representation Tables

This section reports the flexible-model diagnostic table for every representation–task combination. The reported metric is the cross-validated primary metric ($R^2$ for regression, AUROC for classification), and the model column gives the best learner selected within that representation–task scenario. These rows provide the source values for the main-paper family-level comparisons and extended diagnostic context. Representation labels: MolCodon BOW/MolCodon TF-IDF/MolCodon Trace/MolCodon Hybrid/MolCodon Role-Aware/MolCodon Super-Hybrid (MolCodon), SELFIES BOW, SELFIES TF-IDF (SELFIES), Group SELFIES BOW, Group SELFIES TF-IDF (Group SELFIES), ECFP4-2048, FCFP6-2048, RDKit path-2048, MACCS-166 (fingerprints).

Table S5: Flexible-model per-task representation grid.

| Task | Family | Repr. | Best Model | CV main | Test main | Aux 1 | Aux 2 |
|---|---|---|---|---|---|---|---|
| BCL2 Regression | MolCodon | MolCodon BOW | xgboost | 0.848 | 0.857 | 0.926 | 0.761 |
| BCL2 Regression | MolCodon | MolCodon TF-IDF | xgboost | 0.868 | 0.855 | 0.925 | 0.766 |
| BCL2 Regression | MolCodon | MolCodon Trace | xgboost | 0.870 | 0.878 | 0.937 | 0.704 |
| BCL2 Regression | MolCodon | MolCodon Hybrid | xgboost | 0.873 | 0.876 | 0.936 | 0.710 |
| BCL2 Regression | MolCodon | MolCodon Role-Aware | rand. for | 0.861 | 0.856 | 0.926 | 0.763 |
| BCL2 Regression | MolCodon | MolCodon Super-Hybrid | xgboost | 0.872 | 0.877 | 0.937 | 0.706 |
| BCL2 Regression | SELFIES | SELFIES BOW | rand. for | 0.862 | 0.865 | 0.930 | 0.739 |
| BCL2 Regression | SELFIES | SELFIES TF-IDF | xgboost | 0.853 | 0.864 | 0.930 | 0.742 |
| BCL2 Regression | Group SELFIES | Group SELFIES BOW | lgbm | 0.859 | 0.870 | 0.933 | 0.726 |
| BCL2 Regression | Group SELFIES | Group SELFIES TF-IDF | xgboost | 0.870 | 0.892 | 0.945 | 0.662 |
| BCL2 Regression | Fingerprint | ECFP4-2048 | xgboost | 0.887 | 0.906 | 0.952 | 0.618 |
| BCL2 Regression | Fingerprint | FCFP6-2048 | xgboost | 0.880 | 0.908 | 0.953 | 0.609 |
| BCL2 Regression | Fingerprint | RDKit path-2048 | rand. for | 0.878 | 0.914 | 0.957 | 0.592 |
| BCL2 Regression | Fingerprint | MACCS-166 | xgboost | 0.853 | 0.882 | 0.939 | 0.692 |
| HIV Regression | MolCodon | MolCodon BOW | rand. for | 0.573 | 0.612 | 0.783 | 0.917 |
| HIV Regression | MolCodon | MolCodon TF-IDF | lgbm | 0.587 | 0.642 | 0.804 | 0.881 |
| HIV Regression | MolCodon | MolCodon Trace | rand. for | 0.613 | 0.615 | 0.787 | 0.913 |
| HIV Regression | MolCodon | MolCodon Hybrid | lgbm | 0.613 | 0.656 | 0.812 | 0.863 |
| HIV Regression | MolCodon | MolCodon Role-Aware | rand. for | 0.582 | 0.587 | 0.770 | 0.946 |
| HIV Regression | MolCodon | MolCodon Super-Hybrid | lgbm | 0.615 | 0.659 | 0.814 | 0.859 |
| HIV Regression | SELFIES | SELFIES BOW | rand. for | 0.576 | 0.612 | 0.791 | 0.917 |
| HIV Regression | SELFIES | SELFIES TF-IDF | lgbm | 0.575 | 0.617 | 0.790 | 0.911 |
| HIV Regression | Group SELFIES | Group SELFIES BOW | rand. for | 0.568 | 0.598 | 0.778 | 0.933 |
| HIV Regression | Group SELFIES | Group SELFIES TF-IDF | lgbm | 0.573 | 0.630 | 0.797 | 0.895 |
| HIV Regression | Fingerprint | ECFP4-2048 | rand. for | 0.656 | 0.665 | 0.817 | 0.851 |
| HIV Regression | Fingerprint | FCFP6-2048 | rand. for | 0.647 | 0.667 | 0.819 | 0.850 |
| HIV Regression | Fingerprint | RDKit path-2048 | lgbm | 0.676 | 0.685 | 0.828 | 0.826 |
| HIV Regression | Fingerprint | MACCS-166 | rand. for | 0.612 | 0.628 | 0.793 | 0.898 |
| MoleculeNet QM7 | MolCodon | MolCodon BOW | xgboost | 0.478 | 0.462 | 0.680 | 161.7 |
| MoleculeNet QM7 | MolCodon | MolCodon TF-IDF | lgbm | 0.754 | 0.734 | 0.857 | 113.6 |
| MoleculeNet QM7 | MolCodon | MolCodon Trace | lgbm | 0.763 | 0.741 | 0.861 | 112.2 |
| MoleculeNet QM7 | MolCodon | MolCodon Hybrid | lgbm | 0.779 | 0.762 | 0.873 | 107.5 |
| MoleculeNet QM7 | MolCodon | MolCodon Role-Aware | ridge | 0.488 | 0.507 | 0.712 | 154.9 |
| MoleculeNet QM7 | MolCodon | MolCodon Super-Hybrid | lgbm | 0.782 | 0.766 | 0.875 | 106.6 |
| MoleculeNet QM7 | SELFIES | SELFIES BOW | lgbm | 0.440 | 0.442 | 0.665 | 164.7 |
| MoleculeNet QM7 | SELFIES | SELFIES TF-IDF | ridge | 0.643 | 0.616 | 0.785 | 136.7 |
| MoleculeNet QM7 | Group SELFIES | Group SELFIES BOW | lgbm | 0.473 | 0.460 | 0.678 | 162.1 |
| MoleculeNet QM7 | Group SELFIES | Group SELFIES TF-IDF | lgbm | 0.655 | 0.640 | 0.801 | 132.3 |
| MoleculeNet QM7 | Fingerprint | ECFP4-2048 | ridge | 0.538 | 0.502 | 0.710 | 155.6 |
| MoleculeNet QM7 | Fingerprint | FCFP6-2048 | ridge | 0.447 | 0.427 | 0.655 | 167.0 |
| MoleculeNet QM7 | Fingerprint | RDKit path-2048 | lgbm | 0.620 | 0.587 | 0.767 | 141.6 |
| MoleculeNet QM7 | Fingerprint | MACCS-166 | lgbm | 0.604 | 0.572 | 0.757 | 144.2 |

*(continued from previous page)*

| Task | Family | Repr. | Best Model | CV main | Test main | Aux 1 | Aux 2 |
|---|---|---|---|---|---|---|---|
| MoleculeNet ESOL | MolCodon | MolCodon BOW | xgboost | 0.768 | 0.769 | 0.877 | 0.984 |
| MoleculeNet ESOL | MolCodon | MolCodon TF-IDF | lgbm | 0.873 | 0.868 | 0.933 | 0.745 |
| MoleculeNet ESOL | MolCodon | MolCodon Trace | xgboost | 0.853 | 0.870 | 0.934 | 0.738 |
| MoleculeNet ESOL | MolCodon | MolCodon Hybrid | lgbm | 0.874 | 0.879 | 0.939 | 0.713 |
| MoleculeNet ESOL | MolCodon | MolCodon Role-Aware | lgbm | 0.774 | 0.780 | 0.885 | 0.961 |
| MoleculeNet ESOL | MolCodon | MolCodon Super-Hybrid | lgbm | 0.877 | 0.889 | 0.944 | 0.682 |
| MoleculeNet ESOL | SELFIES | SELFIES BOW | xgboost | 0.656 | 0.609 | 0.781 | 1.281 |
| MoleculeNet ESOL | SELFIES | SELFIES TF-IDF | lgbm | 0.777 | 0.796 | 0.893 | 0.926 |
| MoleculeNet ESOL | Group SELFIES | Group SELFIES BOW | xgboost | 0.663 | 0.638 | 0.799 | 1.233 |
| MoleculeNet ESOL | Group SELFIES | Group SELFIES TF-IDF | xgboost | 0.784 | 0.803 | 0.901 | 0.908 |
| MoleculeNet ESOL | Fingerprint | ECFP4-2048 | xgboost | 0.665 | 0.628 | 0.793 | 1.249 |
| MoleculeNet ESOL | Fingerprint | FCFP6-2048 | rand. for | 0.696 | 0.719 | 0.856 | 1.086 |
| MoleculeNet ESOL | Fingerprint | RDKit path-2048 | lgbm | 0.766 | 0.788 | 0.890 | 0.942 |
| MoleculeNet ESOL | Fingerprint | MACCS-166 | xgboost | 0.767 | 0.790 | 0.889 | 0.938 |
| MoleculeNet FreeSolv | MolCodon | MolCodon BOW | xgboost | 0.850 | 0.894 | 0.945 | 1.298 |
| MoleculeNet FreeSolv | MolCodon | MolCodon TF-IDF | xgboost | 0.852 | 0.855 | 0.925 | 1.514 |
| MoleculeNet FreeSolv | MolCodon | MolCodon Trace | xgboost | 0.863 | 0.877 | 0.938 | 1.397 |
| MoleculeNet FreeSolv | MolCodon | MolCodon Hybrid | ridge | 0.867 | 0.899 | 0.949 | 1.264 |
| MoleculeNet FreeSolv | MolCodon | MolCodon Role-Aware | ridge | 0.701 | 0.763 | 0.876 | 1.939 |
| MoleculeNet FreeSolv | MolCodon | MolCodon Super-Hybrid | ridge | 0.868 | 0.901 | 0.950 | 1.253 |
| MoleculeNet FreeSolv | SELFIES | SELFIES BOW | xgboost | 0.732 | 0.846 | 0.921 | 1.564 |
| MoleculeNet FreeSolv | SELFIES | SELFIES TF-IDF | xgboost | 0.779 | 0.825 | 0.909 | 1.666 |
| MoleculeNet FreeSolv | Group SELFIES | Group SELFIES BOW | xgboost | 0.735 | 0.861 | 0.931 | 1.487 |
| MoleculeNet FreeSolv | Group SELFIES | Group SELFIES TF-IDF | lgbm | 0.778 | 0.807 | 0.902 | 1.748 |
| MoleculeNet FreeSolv | Fingerprint | ECFP4-2048 | ridge | 0.789 | 0.764 | 0.878 | 1.933 |
| MoleculeNet FreeSolv | Fingerprint | FCFP6-2048 | xgboost | 0.799 | 0.810 | 0.901 | 1.733 |
| MoleculeNet FreeSolv | Fingerprint | RDKit path-2048 | xgboost | 0.796 | 0.785 | 0.888 | 1.845 |
| MoleculeNet FreeSolv | Fingerprint | MACCS-166 | xgboost | 0.880 | 0.923 | 0.962 | 1.104 |
| MoleculeNet Lipophilicity | MolCodon | MolCodon BOW | lgbm | 0.433 | 0.464 | 0.686 | 0.893 |
| MoleculeNet Lipophilicity | MolCodon | MolCodon TF-IDF | lgbm | 0.537 | 0.547 | 0.745 | 0.820 |
| MoleculeNet Lipophilicity | MolCodon | MolCodon Trace | lgbm | 0.559 | 0.589 | 0.769 | 0.782 |
| MoleculeNet Lipophilicity | MolCodon | MolCodon Hybrid | ridge | 0.581 | 0.575 | 0.758 | 0.795 |
| MoleculeNet Lipophilicity | MolCodon | MolCodon Role-Aware | rand. for | 0.387 | 0.368 | 0.613 | 0.969 |
| MoleculeNet Lipophilicity | MolCodon | MolCodon Super-Hybrid | ridge | 0.595 | 0.594 | 0.771 | 0.777 |
| MoleculeNet Lipophilicity | SELFIES | SELFIES BOW | lgbm | 0.387 | 0.401 | 0.634 | 0.944 |
| MoleculeNet Lipophilicity | SELFIES | SELFIES TF-IDF | lgbm | 0.479 | 0.505 | 0.714 | 0.857 |
| MoleculeNet Lipophilicity | Group SELFIES | Group SELFIES BOW | lgbm | 0.408 | 0.432 | 0.660 | 0.919 |
| MoleculeNet Lipophilicity | Group SELFIES | Group SELFIES TF-IDF | lgbm | 0.503 | 0.564 | 0.756 | 0.805 |
| MoleculeNet Lipophilicity | Fingerprint | ECFP4-2048 | lgbm | 0.555 | 0.544 | 0.740 | 0.824 |
| MoleculeNet Lipophilicity | Fingerprint | FCFP6-2048 | lgbm | 0.552 | 0.567 | 0.756 | 0.802 |
| MoleculeNet Lipophilicity | Fingerprint | RDKit path-2048 | lgbm | 0.463 | 0.490 | 0.705 | 0.870 |
| MoleculeNet Lipophilicity | Fingerprint | MACCS-166 | lgbm | 0.541 | 0.528 | 0.728 | 0.837 |

*(continued from previous page)*

| Task | Family | Repr. | Best Model | CV main | Test main | Aux 1 | Aux 2 |
|---|---|---|---|---|---|---|---|
| MoleculeNet BBBP | MolCodon | MolCodon BOW | xgboost | 0.923 | 0.921 | 0.971 | 0.657 |
| MoleculeNet BBBP | MolCodon | MolCodon TF-IDF | xgboost | 0.922 | 0.916 | 0.969 | 0.668 |
| MoleculeNet BBBP | MolCodon | MolCodon Trace | rand. for | 0.918 | 0.909 | 0.961 | 0.671 |
| MoleculeNet BBBP | MolCodon | MolCodon Hybrid | xgboost | 0.925 | 0.918 | 0.969 | 0.650 |
| MoleculeNet BBBP | MolCodon | MolCodon Role-Aware | rand. for | 0.889 | 0.884 | 0.949 | 0.651 |
| MoleculeNet BBBP | MolCodon | MolCodon Super-Hybrid | rand. for | 0.924 | 0.918 | 0.964 | 0.636 |
| MoleculeNet BBBP | SELFIES | SELFIES BOW | lgbm | 0.909 | 0.870 | 0.945 | 0.543 |
| MoleculeNet BBBP | SELFIES | SELFIES TF-IDF | rand. for | 0.911 | 0.913 | 0.960 | 0.667 |
| MoleculeNet BBBP | Group SELFIES | Group SELFIES BOW | rand. for | 0.903 | 0.903 | 0.961 | 0.629 |
| MoleculeNet BBBP | Group SELFIES | Group SELFIES TF-IDF | rand. for | 0.914 | 0.926 | 0.965 | 0.669 |
| MoleculeNet BBBP | Fingerprint | ECFP4-2048 | rand. for | 0.924 | 0.909 | 0.966 | 0.692 |
| MoleculeNet BBBP | Fingerprint | FCFP6-2048 | rand. for | 0.916 | 0.912 | 0.961 | 0.718 |
| MoleculeNet BBBP | Fingerprint | RDKit path-2048 | xgboost | 0.924 | 0.906 | 0.964 | 0.699 |
| MoleculeNet BBBP | Fingerprint | MACCS-166 | rand. for | 0.922 | 0.931 | 0.975 | 0.688 |
| HIV Classification | MolCodon | MolCodon BOW | rand. for | 0.875 | 0.897 | 0.868 | 0.629 |
| HIV Classification | MolCodon | MolCodon TF-IDF | lgbm | 0.895 | 0.917 | 0.897 | 0.678 |
| HIV Classification | MolCodon | MolCodon Trace | lgbm | 0.901 | 0.917 | 0.896 | 0.696 |
| HIV Classification | MolCodon | MolCodon Hybrid | lgbm | 0.901 | 0.922 | 0.902 | 0.681 |
| HIV Classification | MolCodon | MolCodon Role-Aware | rand. for | 0.892 | 0.908 | 0.879 | 0.645 |
| HIV Classification | MolCodon | MolCodon Super-Hybrid | lgbm | 0.902 | 0.925 | 0.908 | 0.687 |
| HIV Classification | SELFIES | SELFIES BOW | rand. for | 0.889 | 0.896 | 0.860 | 0.651 |
| HIV Classification | SELFIES | SELFIES TF-IDF | lgbm | 0.897 | 0.909 | 0.881 | 0.662 |
| HIV Classification | Group SELFIES | Group SELFIES BOW | lgbm | 0.885 | 0.896 | 0.877 | 0.636 |
| HIV Classification | Group SELFIES | Group SELFIES TF-IDF | lgbm | 0.890 | 0.908 | 0.883 | 0.659 |
| HIV Classification | Fingerprint | ECFP4-2048 | rand. for | 0.914 | 0.930 | 0.914 | 0.720 |
| HIV Classification | Fingerprint | FCFP6-2048 | lgbm | 0.915 | 0.927 | 0.911 | 0.707 |
| HIV Classification | Fingerprint | RDKit path-2048 | xgboost | 0.923 | 0.927 | 0.911 | 0.698 |
| HIV Classification | Fingerprint | MACCS-166 | lgbm | 0.902 | 0.911 | 0.895 | 0.634 |
| Tox21 NR-AR | MolCodon | MolCodon BOW | xgboost | 0.790 | 0.766 | 0.520 | 0.632 |
| Tox21 NR-AR | MolCodon | MolCodon TF-IDF | xgboost | 0.786 | 0.772 | 0.533 | 0.590 |
| Tox21 NR-AR | MolCodon | MolCodon Trace | xgboost | 0.802 | 0.763 | 0.533 | 0.562 |
| Tox21 NR-AR | MolCodon | MolCodon Hybrid | xgboost | 0.790 | 0.771 | 0.524 | 0.603 |
| Tox21 NR-AR | MolCodon | MolCodon Role-Aware | rand. for | 0.813 | 0.773 | 0.460 | 0.524 |
| Tox21 NR-AR | MolCodon | MolCodon Super-Hybrid | xgboost | 0.786 | 0.763 | 0.531 | 0.608 |
| Tox21 NR-AR | SELFIES | SELFIES BOW | xgboost | 0.787 | 0.738 | 0.517 | 0.597 |
| Tox21 NR-AR | SELFIES | SELFIES TF-IDF | rand. for | 0.793 | 0.771 | 0.522 | 0.567 |
| Tox21 NR-AR | Group SELFIES | Group SELFIES BOW | rand. for | 0.799 | 0.749 | 0.468 | 0.593 |
| Tox21 NR-AR | Group SELFIES | Group SELFIES TF-IDF | xgboost | 0.797 | 0.792 | 0.537 | 0.597 |
| Tox21 NR-AR | Fingerprint | ECFP4-2048 | xgboost | 0.803 | 0.770 | 0.524 | 0.608 |
| Tox21 NR-AR | Fingerprint | FCFP6-2048 | rand. for | 0.787 | 0.772 | 0.513 | 0.581 |
| Tox21 NR-AR | Fingerprint | RDKit path-2048 | xgboost | 0.787 | 0.759 | 0.508 | 0.569 |
| Tox21 NR-AR | Fingerprint | MACCS-166 | xgboost | 0.809 | 0.827 | 0.531 | 0.593 |

*(continued from previous page)*

| Task | Family | Repr. | Best Model | CV main | Test main | Aux 1 | Aux 2 |
|---|---|---|---|---|---|---|---|
| Tox21 SR-p53 | MolCodon | MolCodon BOW | rand. for | 0.833 | 0.841 | 0.334 | 0.319 |
| Tox21 SR-p53 | MolCodon | MolCodon TF-IDF | lgbm | 0.849 | 0.840 | 0.383 | 0.299 |
| Tox21 SR-p53 | MolCodon | MolCodon Trace | rand. for | 0.832 | 0.820 | 0.362 | 0.319 |
| Tox21 SR-p53 | MolCodon | MolCodon Hybrid | xgboost | 0.850 | 0.835 | 0.391 | 0.303 |
| Tox21 SR-p53 | MolCodon | MolCodon Role-Aware | rand. for | 0.807 | 0.807 | 0.295 | 0.285 |
| Tox21 SR-p53 | MolCodon | MolCodon Super-Hybrid | xgboost | 0.851 | 0.838 | 0.395 | 0.316 |
| Tox21 SR-p53 | SELFIES | SELFIES BOW | rand. for | 0.811 | 0.795 | 0.291 | 0.270 |
| Tox21 SR-p53 | SELFIES | SELFIES TF-IDF | rand. for | 0.811 | 0.821 | 0.333 | 0.326 |
| Tox21 SR-p53 | Group SELFIES | Group SELFIES BOW | rand. for | 0.807 | 0.815 | 0.303 | 0.314 |
| Tox21 SR-p53 | Group SELFIES | Group SELFIES TF-IDF | rand. for | 0.820 | 0.813 | 0.318 | 0.340 |
| Tox21 SR-p53 | Fingerprint | ECFP4-2048 | rand. for | 0.838 | 0.822 | 0.411 | 0.305 |
| Tox21 SR-p53 | Fingerprint | FCFP6-2048 | rand. for | 0.822 | 0.841 | 0.408 | 0.334 |
| Tox21 SR-p53 | Fingerprint | RDKit path-2048 | rand. for | 0.828 | 0.846 | 0.375 | 0.361 |
| Tox21 SR-p53 | Fingerprint | MACCS-166 | rand. for | 0.848 | 0.841 | 0.349 | 0.363 |
